# Changes in Acceptance of Evolution and Associated Factors during a Year of Introductory Biology: The Shifting Impacts of Biology Knowledge, Politics, Religion, Demographics, and Understandings of the Nature of Science

**DOI:** 10.1101/280479

**Authors:** Ryan D.P. Dunk, Jason R. Wiles

## Abstract

Recent research has identified many factors influencing student acceptance of biological evolution, but few of these factors have been measured in a longitudinal context of changing knowledge and acceptance of evolution over a period of instruction. This study investigates factors previously associated with evolution acceptance as well as other potential factors among students over the course of a year-long majors and non-majors introductory biology sequence at a private, research-intensive university in the northeastern United States. Acceptance of evolution was measured using the Measure of Acceptance of the Theory of Evolution (MATE) instrument, and other factors were measured using well-established instruments and a demographic survey. As expected given the context, evolution was widely accepted among the population (71% of our sample scored in the “high” or “very high” acceptance range), but 160 students were in the very low to moderate acceptance range. Over the course of the academic year, regressions on measures of normalized change revealed that as knowledge of the Nature of Science (NOS) increased, evolution acceptance increased (*R*^2^ = .378, *p* << 0.001). Increasing levels of genetic literacy (*R*^2^ = .214, *p* << 0.001) and Evolutionary Knowledge (*R*^2^ = .177, *p* << 0.001) were also significantly associated with increases in acceptance of evolution. We also examined the longitudinal effect of combining various factors into unified working models of acceptance of evolution, and this is the first study by our knowledge to do so. From fall to spring, the influence of student knowledge of NOS on evolution acceptance increased, as did the influence of genetic literacy. Conversely, the influence of religious variables decreased, as did the influence of political inclinations and race/ethnicity. Our results indicate that as students learn more about the nature of science, they may rely more on scientific explanations for natural phenomena. This study also underscores the importance of using longitudinal, multifactorial analyses to understand acceptance of evolution.

## Introduction

Evolution is the unifying theme of all biology, though which living organisms and communities can be understood most clearly (Dobzhansky, 1973). This framework for the life sciences is reflected in the overwhelming acceptance of evolution amongst biologists (Graffin, 2003). However, acceptance of evolution is not nearly as universal amongst members of the general public as it is in the scientific community. Despite decades of reform to improve evolutionary understanding, in the United States little change has been seen in the number of people who accept evolutionary explanations of life’s diversity as compared to supernatural ones (Gallup, 2014).

Rejection of evolution and the theory around it leads to an inability to understand and to reason about biology as it is studied, understood, and applied by working biologists (Dobzhansky, 1973). The ubiquity of evolutionary theory in the practice of biology makes it challenging to fully understand or engage in biological investigation without a thorough understanding of evolution. Thus, full participation in biology is hindered by a student’s rejection of evolution as a guiding principle of the field. If students are to be well prepared to understand the natural sciences, they should be well educated in evolutionary theory, with attention paid to practices that might mitigate the cognitive barrier of evolution rejection.

Understanding and earnest consideration of evolution is an important goal for non-scientists as well. Evolutionary principles underlie public health issues including vaccinations, antibiotic resistance, and epidemiology; ecological concerns such as invasive species, the biological impacts of climate change, and other environmental implications of human activity; and food security such as pesticide resistance, food crop diversity, and agricultural practices in light of a changing global climate. In addition, science denial by those responsible for guiding public policy may lead to ill-informed decisions and poor potential outcomes regarding future funding for biological sciences. It is for these reasons and more that a general public knowledgeable about evolutionary biology and aware and supportive of its central role in the life sciences is not only desirable, but necessary.

A recent review by Pobiner (2016) includes a thorough overview of the state of evolution acceptance research, and we refer it as a resource to readers who seek an extensive background to the current understanding of factors related to evolution acceptance. Here, we briefly summarize the key factors known to affect evolution acceptance.

Knowledge of evolution is perhaps one of the most intuitive factors related to evolution acceptance; multiple studies have found that a significant positive relationship exists between evolution acceptance and evolutionary knowledge (Brown, 2015; Carter & Wiles, 2014; Deniz, Donnelly, & Yilmaz, 2008; Dorner, 2016; Glaze, Goldston, & Dantzler, 2015; Manwaring, Jensen, Gill, & Bybee, 2015; Rutledge & Warden, 1999). However, this relationship tends to be weaker than would be expected if knowledge was the only (or even the main) factor affecting acceptance of evolution in US populations. Other authors have found no significant relationship: Sinatra et al. found a significant correlation between acceptance of photosynthesis and photosynthesis knowledge while evolution knowledge and acceptance had no such correlation (Sinatra, Southerland, McConaughy, & Demastes, 2003). Similarly,Cavallo and McCall (2008) found no significant impact of evolutionary knowledge on acceptance of evolution, but found that beliefs about the nature of science did have a significant impact.

An understanding of the nature of science has been much more consistently linked to evolution acceptance, with over three decades of results indicating that understanding the aims, processes, and limitations of scientific knowledge leads to an improved acceptance of evolution (Akyol, Tekkaya, Sungur, & Traynor, 2012; Carter & Wiles, 2014; Cavallo & McCall, 2008; Dorner, 2016; Glaze et al., 2015; Johnson & Peeples, 1987; Lombrozo, Thanukos, & Weisberg, 2008; Rutledge & Mitchell, 2002; Trani, 2004). Aside from the overwhelming direct evidence, support for the importance of nature of science in evolution acceptance also comes from an overview of creationist arguments against evolution, which often display fundamental misunderstandings of the nature of science (Eldredge, 2000; Matthews, 1997; Pigliucci, 2008).

Beyond direct creationist rhetoric and understandings, religious affiliation and degree of religiosity also have been shown to affect attitudes towards evolution. While certain denominations outwardly reject evolutionary biology (“Resolution on Scientific Creationism,” 1982), many are more supportive or accommodating of evolutionary ideas (“The Clergy Letter Project,” 2004). However, regardless of the official stance of an individual’s denomination, there is a greater cultural belief among many that evolution and religion are necessarily in conflict (Meadows, Doster, & Jackson, 2000). This commonly held dichotomy is often not addressed by biology instructors who do not discuss religious concerns when presenting evolution in their classrooms (Barnes & Brownell, 2016). This might lead to an understanding of religious experience as standing in opposition to scientific exploration, and sets up intensity of religious belief (or “religiosity”) as a more direct way to test the relationship between religion and evolution acceptance. Many studies have done so, and have found that increased religiosity is associated with decreased acceptance of evolution (Brown, 2015; Carter & Wiles, 2014; Glaze et al., 2015; Heddy & Nadelson, 2013; Lombrozo et al., 2008; Manwaring et al., 2015; Moore, Brooks, & Cotner, 2011; Nadelson & Hardy, 2015; Rissler, Duncan, & Caruso, 2014; Trani, 2004). Religiosity, however, is a complicated construct (P. C. Hill & Hood, 1999), referring to both *intrinsic* religiosity (the degree to which religion influences personal understanding and decision making) and *extrinsic* religiosity (the importance of religious worship and religious communities for an individual). For the remainder of this article we will consider only intrinsic religiosity.

Acceptance of evolution is also impacted by political ideology. People in the United States who identify as Republican or as conservative tend to reject evolution as an explanation for human life on earth at a greater rate than their more centrist and liberal peers (Newport, 2007; Pew Research Center, 2015). This trend was also found to be significant in studies that used multifactorial models from large survey data (Baker, 2013; Mazur, 2004) and those that looked specifically at acceptance of evolution in university students (Carter & Wiles, 2014; Cotner, Brooks, & Moore, 2014; Hawley, Short, McCune, Osman, & Little, 2011; Nadelson & Hardy, 2015).

A number of various, but related, psychological factors have also been found to impact evolution acceptance. Thinking dispositions such as Actively Open-Minded Thinking (openness to ideas that conflict with one’s own) have been found to significantly impact evolution acceptance (Deniz et al., 2008; Sinatra et al., 2003). Sinatra et al. (2003) also found students’ levels of epistemological sophistication (the tendency to rely on authority and view knowledge in absolute terms) to be significantly related to evolution acceptance. Finally, other authors have found openness to experience, one of the “Big Five” personality traits that measures intellectualism and creativity (John, Naumann, & Soto, 2008) to be significantly related to acceptance of evolution as well (Hawley et al., 2011).

A host of other variables, which we will for convenience refer to under the umbrella term of “demographic variables”, have been found to be significantly related to acceptance of evolution. Of most relevance to the current study, different researchers have found age (Gallup, 2014; Mazur, 2004), sex and gender (Baker, 2013; Grose & Simpson, 1982; Miller, Scott, & Okamoto, 2006), academic major (Flower, 2006; Ha, Cha, & Ku, 2012), geographic location (Mazur, 2004; Miller et al., 2006), rurality (Baker, 2013; Mazur, 2004), youth science exposure (Hawley et al., 2011; Short & Hawley, 2012), interest in science (Ha et al., 2012; Lombrozo et al., 2008), level of biology preparation (Lord & Marino, 1993; Rice, Olson, & Colbert, 2011), parents’ level of education (Hawley et al., 2011), and number of religious friends (J. P. Hill, 2014) to be significantly associated with evolution acceptance. Race and ethnicity is another key demographic variable of interest since in the United States race is an extremely salient factor in educational access and experience (Howard & Navarro, 2016; Ladson-Billinngs & Tate, 1995). Though previous research has tended to find no significant relationship between race or ethnicity and evolution acceptance (Dorner, 2016; Nadelson & Hardy, 2015; Woods & Scharmann, 2001), we feel it is important to include and continue to study, especially in light of Walls’ (2016, p.1) challenge for racially inclusive science education: “science education research aimed at improving an individual’s science learning and understanding necessarily must take into account the background and experiences that could impact the success of such an undertaking.”

Our prior work was among the first studies to combine most of these factors into a single working model (Dunk, Petto, Wiles, & Campbell, 2017). In a midwestern public university setting, we found student understanding of the nature of science to be the most significant factor in our model, explaining over 13% of the unique variation in acceptance of evolution. This was followed in explanatory power by religiosity (10%), openness to experience (5%), knowledge of evolution (3%) and religious denomination (3%). Overall, our model explained over 33% of the variation in our measure of acceptance of evolution, which is quite substantial for a model of human cognition and attitudes.

Here, we investigate the role of factors previously deemed important for acceptance of evolution as well as investigate other potential factors. Specifically, we have extended the theoretical model by applying it to a longitudinal study to measure changes in these variables over time. Prior research (including much of our own) has often been limited in time, presenting a single snapshot of individuals’ acceptance of evolution. However, acceptance of evolution is a construct in flux for many students, attested to by the volumes dedicated to changing acceptance of evolution (via evolution instruction) geared towards instructors (Alters & Alters, 2001; Lynn, Glaze, Evans, & Reed, 2017) or towards the general public (Coyne, 2009; Mayr, 2001; Shermer, 2006). Thus, to better understand the changing nature of evolution acceptance, we conducted the following study to investigate how evolution acceptance and its associated factors may change over time. A longitudinal study allows us to support causal inferences in our models by establishing the associated factors’ continuing or changing relationships with acceptance of evolution.

### Predictions

This study seeks to test two general hypotheses: (i) that when certain variables shown to be related to acceptance of evolution change over time, that change is correlated with change in acceptance of evolution, and (ii) that the amount of variance in acceptance of evolution explained by these variables changes as students progress in knowledge and experience. Specifically, given the previous significant impact on evolution acceptance demonstrated by an understanding of the nature of science, religiosity, openness to experience, and measures of knowledge of evolution (Dunk et al., 2017), we expected to find that changes in these variables would be significantly correlated with changes in evolution acceptance. We expected the direction of these relationships to be positive for nature of science understanding, evolution knowledge measures, and openness to experience (individuals who increase in these variables over time will tend to increase in acceptance of evolution) and negative for intrinsic religiosity (individuals who increase in their intrinsic religiosity will tend to decrease in acceptance of evolution).

Due to the large models employed, along with the paucity of research using multifactorial models on many of the measures employed, it was difficult to make highly specific predictions. However, we were able to make some discrete predictions about the changing influence on evolution acceptance of general groups of variables between the beginning and end of a year of university-level introductory biology instruction. Firstly, we expected that a year of instruction in biology would tend to diminish the effects of prior preparation on evolution acceptance. We believed that this would be most prominent in variables that measure knowledge of evolution or biology either directly or indirectly, but would also extend to more general demographic variables inasmuch as those variables represent differential access to opportunity to engage with evolutionary biology content. Secondly, we expected to find that as students learned more about evolutionary biology, they would tend to rely more on scientific explanations of evolution and other biological phenomena and less on non-scientific (e.g., religious) explanations. This would be measured over the year as a decreased impact of religious variables on acceptance of evolution, and an increased impact of variables related to understanding of the nature of science. Thirdly, we expected that for some, the year in a university setting would provide students with exposure to new ideas, philosophies, and personalities. Thus, we expected that the levels of an individual’s openness to experience would become more important as the year progressed. This would also be reflected in a decreased importance of political views and political party affiliation on acceptance of evolution, as students who may have been surrounded by more conservative social environments that tend to be less tolerant of evolutionary ideas were exposed to ideas in counterpoint throughout the year of biology instruction and other aspects of the university experience.

## Methods

### Data Collection

Introductory biology students (*N* = 656) at a private northeastern university were surveyed under an IRB approved protocol at the beginning and end of a year-long biology course. The introductory biology course is a survey course required for biology majors and majors in related disciplines, but also popular among non-majors for fulfilling general education requirements. The full course is composed of a two-semester (Fall-Spring) sequence, though it is sometimes (rarely) taken out of sequence by some students. Completion of the sequence is not mandatory for all students, but most students take both semesters. Surveys were administered online through course management software tools (Blackboard) at the beginning of the fall and end of the spring semesters (hereafter, “fall” and “spring”). Participation was voluntary, and students received a small amount of extra credit for participation (1 point out of 1,000 per survey instrument). The survey consisted of 6 different instruments, with a 7th survey asking for participants’ demographic information, for a total of 171 individual response items. These surveys are summarized in table 1.

**Table 1.** Surveys used the current study.

| **Survey Coverage** | **Survey Name** | **Citation** |
| --- | --- | --- |
| Acceptance of Evolution | Measure of Acceptance of the Theory of Evolution (MATE) | Rutledge & Sadler, 2007 |
| Knowledge of the Nature of Science | Nature of Scientific Knowledge Survey (NSKS) | Rubba, 1977 |
| Religiosity | Combined version of the Duke University Religion Index (DUREL) and Hoge’s Intrinsic Religious Motivation Scale | Hoge, 1972; Koenig & Büssing, 2010 |
| Epistemological Sophistication | Openness to Experience factor of Big Five Inventory | John et al., 2008 |
| Evolution Knowledge | Genetic Literacy, Evolutionary Knowledge, and Evolutionary Misconceptions factors from Evolutionary Attitudes and Literacy Survey- Short Form (EALS-SF) | Short and Hawley, 2012 |
| Friend Network | Edited portion of National Study of Youth and Religion | Hill, 2014 |
| Demographics | Various studies |  |

Acceptance of evolution, the outcome variable of interest, was measured by the Measure of Acceptance of Evolution (MATE; Rutledge & Sadler, 2007; Rutledge & Warden, 1999). While there are a number of more recent evolutionary acceptance measures (Nadelson & Southerland, 2012; Smith, Snyder, & Devereaux, 2016), the MATE was chosen as it is a consistently valid instrument that allows a comparison between the present study and the many former studies that used the measure previously. Additionally, we are aware of a recent study that finds a potential two-factor structure in the MATE (Romine, Walter, Bosse, & Todd, 2017). However, to explore this in our own data we would want a sample with a larger diversity of academic majors than the current study allows. We utilized the instrument as a single measure, which is a technique that continues to be endorsed by the authors of the two-factor study.

Another survey instrument that deserves special attention is our measure of an individual’s understanding of the aims, processes, and philosophy of science, which are summed up in the term “nature of science”. One of the more popular nature of science scales, the Views of Nature of Science questionnaire (VNOS; Lederman, Abd-El-Khalick, Bell, & Schwartz, 2002), was not used, although we acknowledge its value for providing a rich understanding of individual students’ conceptions of science. The open-ended nature of the VNOS questions and the more qualitative data they return were not suitable for this study. Among the other nature of science scales (many of which are summarized in Lederman, Wade, & Bell, 1998), we chose the Nature of Scientific Knowledge Survey (NSKS; Rubba & Andersen, 1978), a 48-item, 5-point Likert survey tool. Though it has been some time since its original construction, the NSKS is still being used currently (Ozdemir & Dikici, 2017), and has been successful enough to have been translated into multiple languages since its inception (Chan, 2005; Folmer, Barbosa, Soares, & Rocha, 2009; Kilic, Sungur, Cakiroglu, & Tekkaya, 2005).

The NSKS was considered especially beneficial for this study for its dissection of the nature of science into six distinct factors, each separately measurable within the one instrument. The separate factors are defined as follows (with a brief description of each given parenthetically, paraphrased from Rubba & Andersen, 1978): Amoral (scientific knowledge itself cannot be judged as morally right or wrong, although its methods and applications can), Creative (scientific inquiry is a process that relies on creative input from researchers), Developmental (scientific knowledge is not absolute, and subject to change based on additional evidence), Parsimonious (scientific explanations should be as simple and comprehensive as possible), Testable (scientific explanations are capable of being tested and are open to testing and retesting), and Unified (different branches of scientific inquiry allow for specialization, but all science contributes to a single body of mutually intelligible and relevant knowledge). These distinctly measurable factors allow for a more nuanced analysis of changes in the understanding of science, as well as the relationship between the nature of science and acceptance of evolution.

All survey instruments described in Table 1 are 5-item Likert surveys except the factors from the short form of the Evolutionary Attitudes and Literacy Survey (EALS-SF; Short & Hawley, 2012), which are 7-item Likert surveys, and the demographic variables, which vary in form. The demographic questions addressed included gender identity, age, major, race/ethnicity, state or country of origin, rurality of childhood home, childhood informal science exposure, general interest in science, parents’ level of education, religious affiliation/ denomination, level of religious activity, political leanings, and political party affiliation. Parents’ combined level of education was determined by converting each of the ordinal responses to questions on each parents’ education level to a number (1-8, 1 indicating “never attended school or only attended kindergarten” and 8 indicating “post-bachelor’s degree [graduate school, law school, medical school]”) and averaging them; this variable could assume 15 different levels, from 1 to 8 in 0.5 increments, and was thus treated as a continuous rather than categorical variable. Specific wording for the demographic questions can be found in supplemental table S1.

Survey responses were cleaned by invalidating responses that indicated extremely contradictory positions, which was indication of respondent apathy. Additionally, individuals who were under the age of 18 were excluded from research participation. Gender, major, race/ethnicity, census region of origin, and religious affiliation were all coded. Categories in any variable with less than 3% of total responses were dropped (responses nulled); participants with responses indicated as “other” in codes for religion and political party were also removed, as these were a heterogeneous group with results that would not represent an interpretable pattern.

#### Analysis

Summary statistics for all variables were determined from survey responses from the beginning of the fall semester. These allow a description of the survey population as well as an understanding of the baseline values for each of the variables of interest in the study.

Survey response scores from the beginning of the fall semester and the end of the spring semester (representing a year of introductory biology education) were compared using normalized change (Marx & Cummings, 2007), a metric of change or improvement that attempts to eliminate both ceiling effects and pre-test score bias. Normalized change is similar to normalized gain and runs from −1 (maximal decrease) to +1 (maximal increase). Normalized change scores for measures of evolutionary knowledge, genetic literacy, evolutionary misconceptions, religiosity, openness to experience, and the 7 measures of knowledge of the nature of science (total score and 6 subscores) were correlated individually to the normalized change scores for acceptance of evolution. P-values for these tests were adjusted for multiple comparison using the Holm-Bonferroni sequential procedure (Abdi, 2010).

To investigate the unique impact of each dependent variable on MATE score in both the fall and spring, multifactorial General Linear Models (GLM) (Huitema, 2011; Rutherford, 2001) were generated for the pre-course and post-course data in a manual stepwise regression fashion. First, individual regressions or one-factor ANOVAs between acceptance of evolution and all other variables in the study were conducted. In total, 16 regressions were conducted (Intrinsic Religiosity, Openness to Experience, NSKS total and all 6 subscales of nature of science conceptions, Evolutionary Misconceptions, Evolutionary Knowledge, Genetic Literacy, age, number of science classes taken in college, number of biology classes taken in college, and parents’ combined level of education) and 18 one-factor ANOVAs were conducted (gender, pre-med status, major or intended major, race/ ethnicity, census region of origin, rurality of childhood home, childhood exposure to science in informal settings, general interest in science, mother’s education level, father’s education level, religious affiliation/ denomination, level of religious activity, general political views, political views on social issues, political views on fiscal issues, political party affiliation, number of religious friends, and number of friends with a similar religion to respondent’s).

Those variables that had a significant (α=0.05) relationship with acceptance of evolution were included as dependent variables into a large multifactorial main effects GLM (the “full model”) with MATE score as the dependent variable. Factors in that model that retained a relationship with acceptance of evolution at an alpha of 0.5 or below were included in the next model. This liberal cutoff level was chosen to ensure that all potentially significant variables were included in the final model. The second model (hereafter, “intermediate model”) was run similarly to the full model, and again variables with an alpha of 0.5 or below were selected to be included in the “final model”. Essentially, iterative models were run until no factors in the model had an alpha above 0.5; this was done with the intent to allow the most power to detect significance levels of the remaining variables in the model. The final model was run as a main effects GLM with acceptance of evolution (as measured by MATE score) as the dependent variable, and the remaining independent variables run as factors (for categorical variables) or covariates (for continuous variables).

This iterative procedure was conducted independently for the data gathered from the beginning of the fall semester and the end of the spring semester. To confirm the differences between the models were due to changes throughout the year and not participant selection, all variables in the fall data set were analyzed for a significant difference between those individuals who went on to the spring semester and those who did not, and all variables in the spring data set were analyzed for a significant difference between those individuals who were enrolled in the fall semester and those who were not. The tests were conducted either as one-factor ANOVAs (for continuous variables) or chi-square tests of independence (for categorical variables). Students who were enrolled in both semesters and students who were enrolled for one semester did not differ for any variables that were included in the main effects GLM after Bonferroni correction for multiple tests.

The main effects models for fall and spring were compared for differences in the structure of the model as well as differences in the overall and relative effect size of each variable in the model. Multicollinearity in the final models was assessed using generalized variance inflation factors (Fox & Monette, 1992) and was found to be within an acceptable limit (all gVIFs were under 2). Effect size (as eta-squared, η^2^; Richardson, 2011) for each variable and P-value adjustments for multiple tests were calculated manually; all other statistical procedures were done in RStudio 1.0.153 (RStudio Team, 2016) running R 3.4.1 (R Core Team, 2017).

## Results

### (i) Descriptive Statistics

Descriptive statistics were calculated for all variables in the fall survey administration. Table 2 shows summary statistics for continuous variables, including mean, maximum, minimum, and standard deviation. Frequency tables for select categorical variables are given in table 3, and frequency tables for all other variables are given in supplemental table S2.

**Table 2.** Summary statistics for continuous variables in the fall survey administration.

|  | **Mean** | **SD** | **Minimum**<br>**(Min Possible)** | **Maximum**<br>**(Max Possible)** |
| --- | --- | --- | --- | --- |
| MATE | 81.00 | 9.66 | 32 (20) | 100 (100) |
| Intrinsic Religiosity | 23.88 | 8.30 | 10 (10) | 50 (50) |
| Openness to Experience | 35.86 | 5.90 | 19 (10) | 49 (50) |
| NSKS Total | 171.68 | 12.24 | 133 (48) | 216 (240) |
| NSKS Amoral | 26.92 | 4.18 | 16 (8) | 38 (40) |
| NSKS Creative | 27.46 | 4.84 | 8 (8) | 40 (40) |
| NSKS Developmental | 30.39 | 3.19 | 20 (8) | 40 (40) |
| NSKS Parsimonious | 22.96 | 3.23 | 14 (8) | 35 (40) |
| NSKS Testable | 31.85 | 3.85 | 19 (8) | 40 (40) |
| NSKS Unified | 31.94 | 3.43 | 20 (8) | 40 (40) |
| Evolutionary Misconceptions | 12.54 | 3.26 | 3 (3) | 21 (21) |
| Evolutionary Knowledge | 26.98 | 3.69 | 16 (5) | 35 (35) |
| Genetic Literacy | 19.97 | 3.60 | 11 (4) | 28 (28) |
| Age | 18.81 | 2.62 | 18 (18) | 64 (∞) |
| No. College Science Classes | 1.56 | 2.06 | 0 (0) | 20 (∞) |
| No. College Biology Classes | 0.25 | 0.66 | 0 (0) | 7 (∞) |
| Parents' Combined Education Level | 6.39 | 1.40 | 2 (0) | 8 (8) |

**Table 3.**
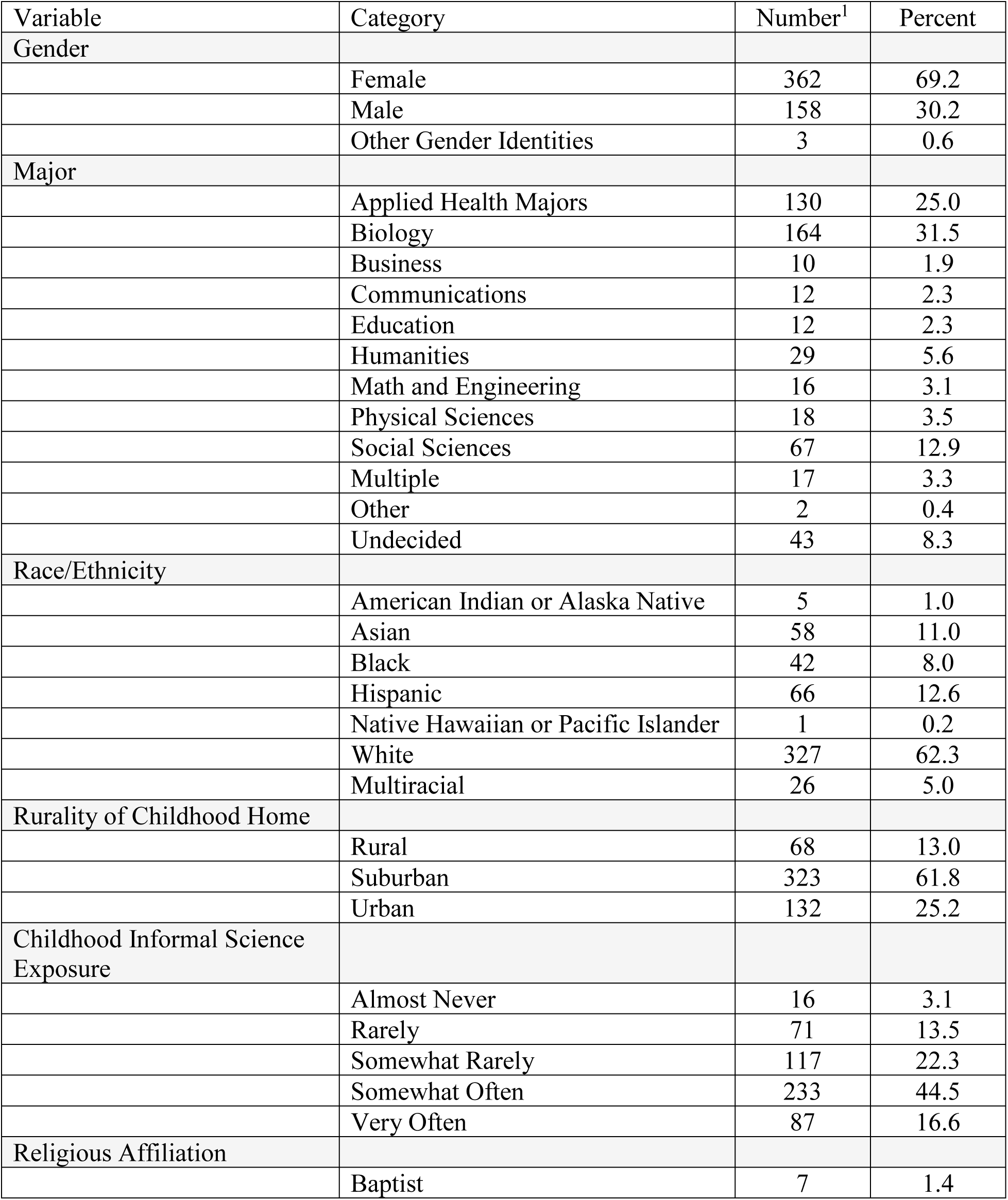
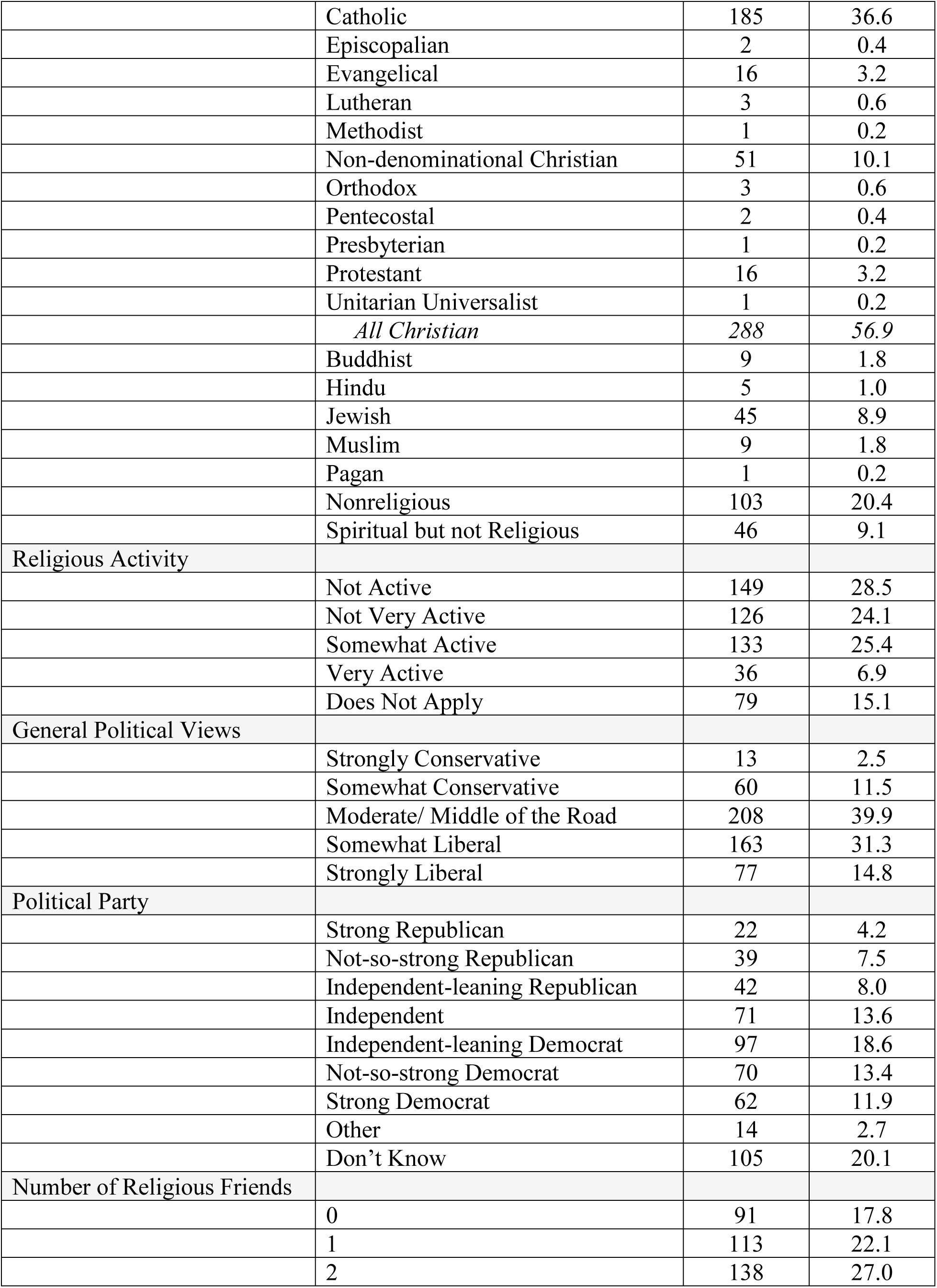
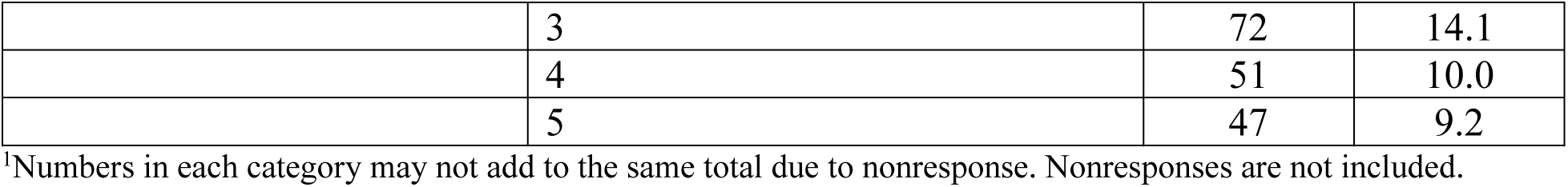
Frequency tables for select categorical variables in the fall survey administration.

The student population in this intro biology class tends to be young (*M* = 18.8, *SD* = 2.6), with a majority (62%) identifying as white. Women were also in the majority (69%). Over a quarter (26%) of the students in the sample identified with racial or ethnic identities that are considered underrepresented in the natural sciences (Snyder, Sloane, Dunk, & Wiles, 2016). There is even greater diversity amongst the population studied in political views, religious affiliations, and other demographics such as childhood exposure to informal science learning.

Cronbach’s alpha was calculated for the dependent variable, MATE score, and was found to be high (0.9). Looking at levels of evolution acceptance, even upon entering the introductory biology course, students’ acceptance of evolution tended to be high (MATE score *M* = 81.0 *SD* = 9.7; table 4). However, a large number of individuals fell into the moderate category, indicating a substantial potential for change among these students toward higher acceptance of evolution. Students’ understanding of the nature of science, evolutionary knowledge, and genetic literacy tended to be more in the middle of the potential range (table 2).

**Table 4.** Levels of evolution acceptance for introductory biology students at the beginning of the fall semester.

| Acceptance level<sup>1</sup> | Score range | Number of respondents |
| --- | --- | --- |
| Very low | 20-52 | 4 |
| Low | 53-64 | 17 |
| Moderate | 65-76 | 118 |
| High | 77-88 | 237 |
| Very high | 89-100 | 108 |

<sup>1</sup>Score range for acceptance levels defined by Rutledge and Sadler (2007).

### (ii) Normalized Change

Normalized change scores for acceptance of evolution were found to be significantly correlated with change in almost all tested associated variables (table 5, figures 1 & 2). The correlation was highest between change in the full nature of science understanding measure and change in acceptance of evolution, although two of the NSKS subscales (Parsimonious and Creative) did not significantly change along with acceptance of evolution. The other four NSKS measures showed a fairly robust relationship in their change throughout the year with acceptance of evolution (figure 2), as did the genetic literacy and evolutionary knowledge factors from the EALS-SF (Short & Hawley, 2012). Normalized change scores in the evolutionary misconceptions factor from the EALS-SF, as well as openness to experience and intrinsic religiosity, had a very modest but still significant relationship with change in acceptance of evolution across the year (figure 1).

**Figure 1.**
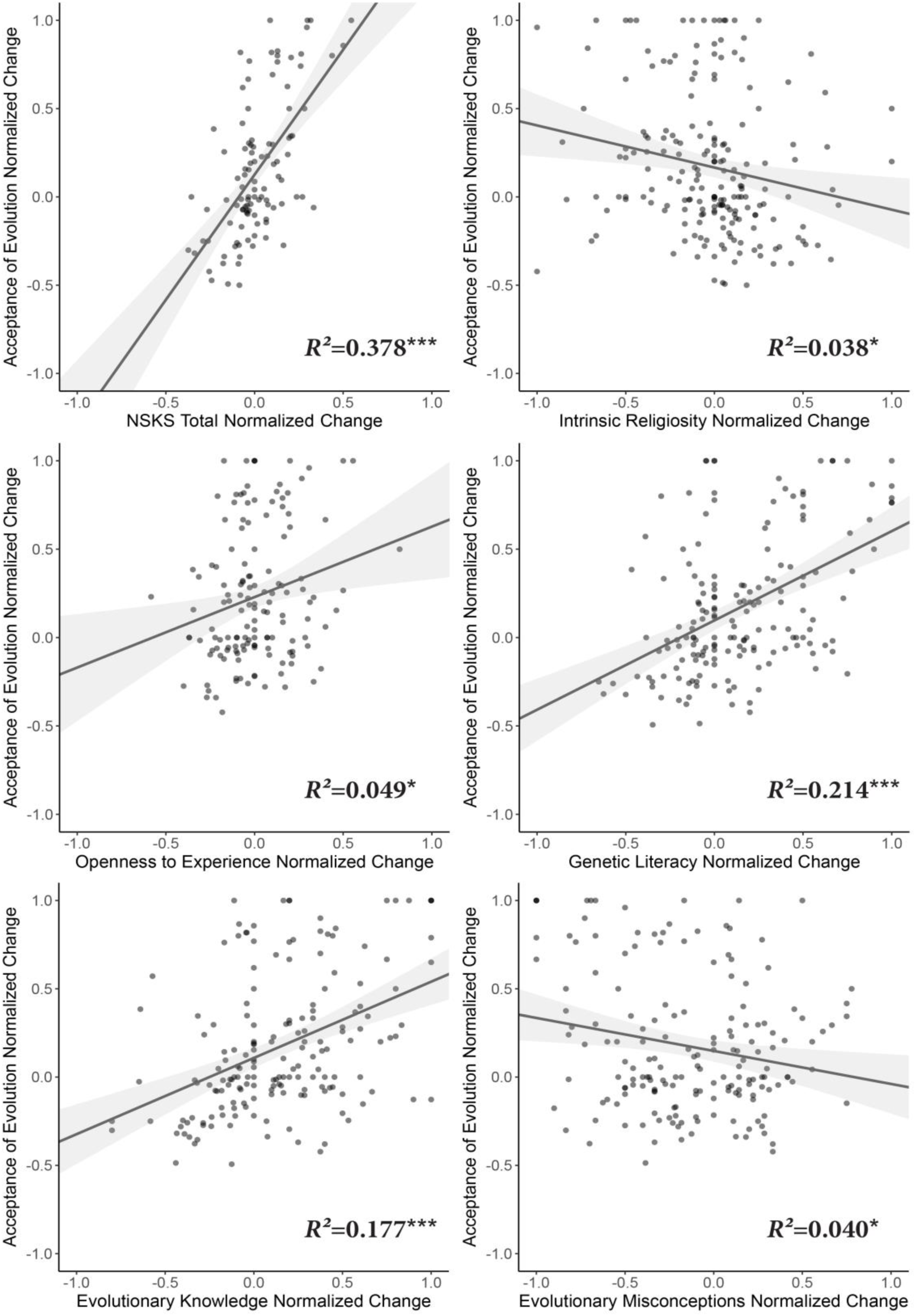
Correlations between normalized change in acceptance of evolution and normalized change in 6 different variables. *R*^2^ values are given on each plot, and shading represents 95% CI of the regression line. Dots are translucent, so darkened areas show overlap of multiple points. Significance: * = *p* < 0.05, ** = *p* < 0.01, *** = *p* < 0.001, ^NS^ = Not Significant.

**Figure 2.**
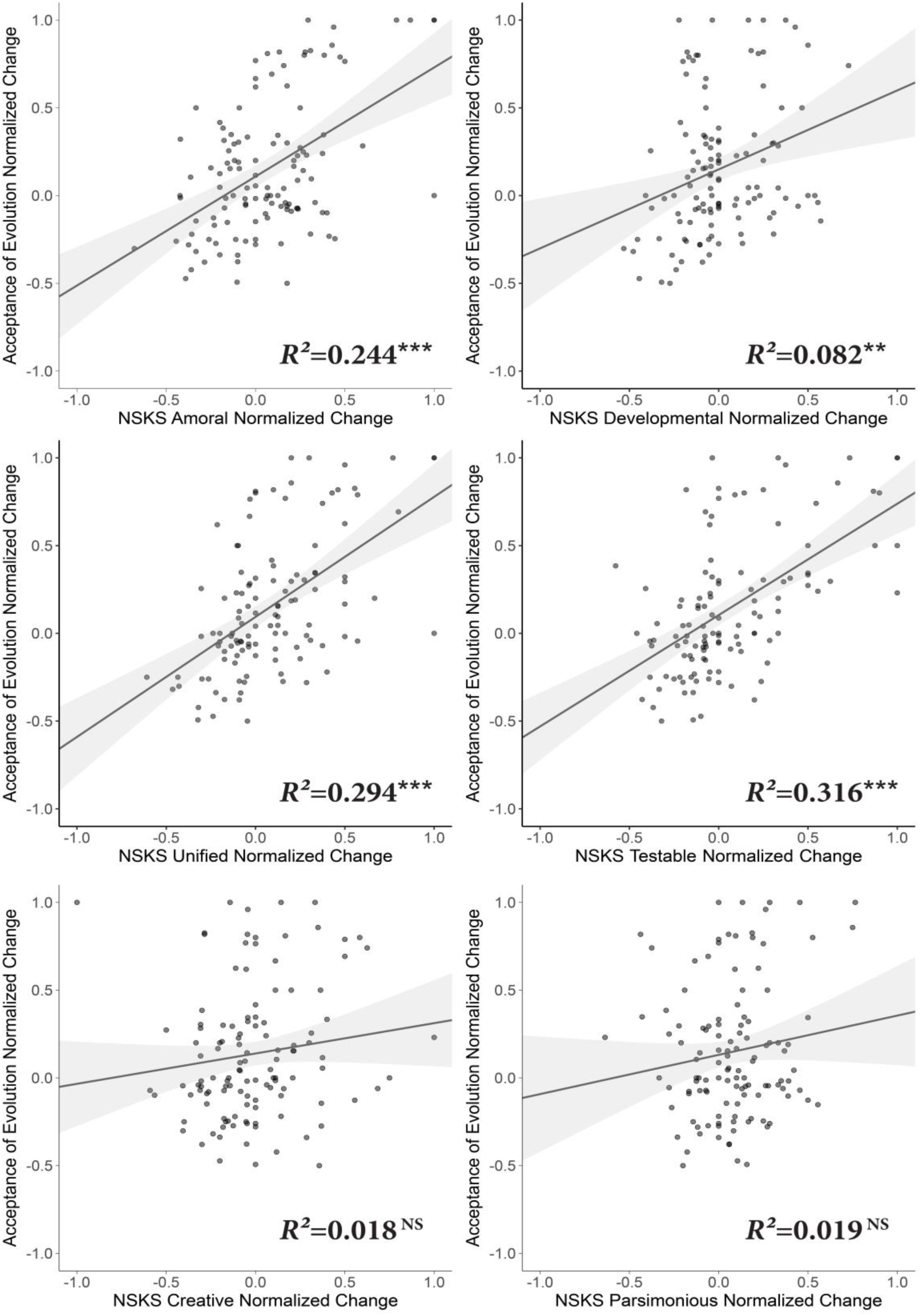
Correlations between normalized change in acceptance of evolution and normalized change in the nature of science variables measured by the NSKS. *R*^2^ values are given on each plot, and shading represents 95% CI of the regression line. Dots are translucent so darkened areas show overlap of multiple points. Significance: * = *p* <0.05, ** = *p* < 0.01, *** = *p* < 0.001, ^NS^ = Not Significant.

**Table 5.** Results of correlations between normalized change of acceptance of evolution (MATE score) and normalized change of 12 different independent variables.

| Variable | $R^2$ | $p_{\text{adj}}^{\dagger}$ |
| --- | --- | --- |
| Nature of Science Understanding (NSKS) | .378 | < .000 001 |
| NSKS Testable | .316 | < .000 001 |
| NSKS Unified | .294 | < .000 001 |
| NSKS Amoral | .244 | < .000 001 |
| NSKS Developmental | .082 | .009 |
| NSKS Parsimonious | .019 | NS |
| NSKS Creative | .018 | NS |
| Genetic Literacy (EALS-SF) | .214 | < .000 001 |
| Evolutionary Knowledge (EALS-SF) | .177 | < .000 001 |
| Evolutionary Misconceptions (EALS-SF) | .040 | .025 |
| Openness to Experience | .049 | .032 |
| Intrinsic Religiosity | .038 | .032 |

<sup>†</sup>Adjusted $p$ values are corrected by Holm-Bonferroni method.

Specifically, we found that a students’ change over the semester in their understanding of the nature of science explained 38% of the change in their acceptance of evolution. This finding was highly significant. Change in evolutionary knowledge was significantly positively associated with change in acceptance of evolution as well (*R*^2^ = 0.17, *p* <0.001). Change in openness to experience had a quite modest relationship with change in acceptance of evolution (*R^2^* = 0.05, *p* = 0.032). Finally, change in intrinsic religiosity had a significant, but quite small, negative relationship with change in acceptance of evolution (*R*^2^ = 0.04, *p* = 0.032).

### (iii) Pre-course and post-course general linear modeling

Data from survey administrations at the beginning of the fall semester and the end of the spring semester were analyzed separately. Individual variable correlation and ANOVA results, as well as the full and intermediate models for both semesters are given in supplemental tables S3–S8. The results of this final model for both semesters are presented in table 6, with variables sorted by general category. Eta-squared (*η*^2^) values are given for comparison both within and between models of each variable’s independent contribution to total differences in acceptance of evolution. Overall, significant terms in the early fall model explained 41% of the total variation in acceptance of evolution, while significant terms in the late spring model explained 39% of the total variation in acceptance of evolution.

**Table 6.** Final general linear models for both the early fall and late spring of a year of introductory biology. Acceptance of evolution (as measured by the MATE) is the dependent variable. (NIFM = not in final model)

| Early Fall |  |  | Late Spring |  |  |  |
| --- | --- | --- | --- | --- | --- | --- |
| Political Variables |  |  |  |  |  |  |
| *F* | *p* | $\eta^2$ | | *F* | *p* | $\eta^2$ |
| 3.44 | 0.002 | .0742 | Political Party | 2.12 | 0.043 | .0411 |
| 3.82 | 0.005 | .0472 | General Political Views | 4.01 | 0.004 | .0444 |
| *Combined $\eta^2$:* |  | .1214 |  | *Combined $\eta^2$:* |  | .0855 |
| Religious Variables |  |  |  |  |  |  |
| *F* | *p* | $\eta^2$ | | *F* | *p* | $\eta^2$ |
| 4.38 | <0.001 | .0810 | Religious Affiliation | 1.48 | 0.177 |  |
| 5.25 | 0.023 | .0162 | Intrinsic Religiosity | 9.01 | 0.003 | .0249 |
| 1.21 | 0.309 |  | Number of Religious Friends | 3.43 | 0.006 | .0474 |
| NIFM |  |  | Religious Activity | 1.04 | 0.390 |  |
| *Combined $\eta^2$:* |  | .0972 |  | *Combined $\eta^2$:* |  | .0723 |
| Nature of Science Variables |  |  |  |  |  |  |
| *F* | *p* | $\eta^2$ | | *F* | *p* | $\eta^2$ |
| 8.36 | 0.004 | .0258 | NSKS Amoral | NIFM |  |  |
| 8.09 | 0.005 | .0249 | NSKS Unified | 15.95 | <0.001 | .0441 |
| NIFM |  |  | NSKS Testable | 15.84 | <0.001 | .0438 |
| NIFM |  |  | NSKS Parsimonious | 0.78 | 0.379 |  |
| *Combined $\eta^2$:* |  | .0508 |  | *Combined $\eta^2$:* |  | .0879 |
| Knowledge Variables |  |  |  |  |  |  |
| *F* | *p* | $\eta^2$ | | *F* | *p* | $\eta^2$ |
| 13.08 | <0.001 | .0403 | Evolutionary Knowledge | 15.28 | <0.001 | .0423 |
| 9.53 | 0.002 | .0294 | Number of College Biology Classes Taken | NIFM |  |  |
| 3.99 | 0.047 | .0123 | Genetic Literacy | 11.34 | <0.001 | .0314 |
| NIFM |  |  | Evolutionary Misconceptions | 0.54 | 0.464 |  |
| *Combined $\eta^2$:* |  | .0820 |  | *Combined $\eta^2$:* |  | .0737 |
| Demographic Variables |  |  |  |  |  |  |
| *F* | *p* | $\eta^2$ | | *F* | *p* | $\eta^2$ |
| 4.34 | 0.002 | .0535 | Race/Ethnicity | 3.20 | 0.014 | .0354 |
| 1.01 | 0.402 |  | Childhood Informal Science Exposure | 3.18 | 0.015 | .0351 |
| 2.35 | 0.099 |  | Rurality | NIFM |  |  |
| *Combined $\eta^2$:* |  | .0535 |  | *Combined $\eta^2$:* |  | .0705 |

## Discussion

### (i) Descriptive Statistics

As noted, the population in our study tends to be young. The majority identify as white, though there is substantial representation from underrepresented racial groups. Women are in the majority. This representation is a common feature of studies that utilize a college undergraduate population, and is very similar to our previous study conducted at a different university (Dunk et al., 2017). Students in this study tended to have a high level of acceptance of evolution at the start of the fall semester, which is also similar to other studies of ours, both at this university (Carter & Wiles, 2014) and elsewhere (Dunk et al., 2017). Although not without precedent in other studies (Dorner & Scott, 2016), MATE scores in this study tended to be higher than other studies that utilize the MATE, regardless of age and experience level of respondents (Cavallo & McCall, 2008; Grossman & Fleet, 2017; Rissler et al., 2014; Rutledge & Sadler, 2007; Wiles & Alters, 2011).

With regard to nature of science conceptions as measured by the NSKS, we found that respondents tended on average to score near the midpoint of the instrument scale on the Amoral, Creative, and Parsimonious factors, but averaged somewhat higher on the Developmental, United, and Testable aspects; this indicates a somewhat higher level of understanding of those factors of the nature of science. Amongst all the factors, it seems that the one least understood by students in this survey was the parsimonious nature of science, as both its mean and its maximum score were the lowest of all the NSKS factors. This is perhaps not surprising, as younger college students tend to view science as complex, and instruction tends to focus on the explanatory power of scientific knowledge, and not its relative simplicity. This pattern of scores, as well as the actual means, closely matches that found by Rubba and Anderson (1978) of non-majors in a biology course in one of the first uses of the NSKS. A somewhat similar pattern is also found in more recent uses of the NSKS (Owens & Foos, 2007), but holds less strongly in international settings (Chan, 2005; Folmer et al., 2009), suggesting the pattern of understanding of the nature of science is not universal and is likely influenced by cultural attitudes and understandings of scientific processes.

### (ii) Normalized Change

Looking at the correlations between normalized change in acceptance of evolution as well as normalized change in the other continuous variables, the strongest relationship was between an understanding of the nature of science and acceptance of evolution. That is, individuals who increased in their understanding of the nature of science were likely to increase in their acceptance of evolution. This relationship was especially strong and significant for the Amoral, Unified, and Testable subscales of the NSKS. Thus, these areas of nature of science might be particularly fruitful towards developing curricular interventions that would lead to both improved understanding of the nature of science and increased evolution acceptance.

Change in openness to experience, as mentioned above, had a comparatively small relationship with change in acceptance of evolution. Though it was found to be significant, the percent of variance explained was much smaller than that for many of the NSKS and EALS-SF variables, indicating that openness to experience may not be a good target for ways to improve evolution acceptance. This is a relatively surprising finding, given the comparatively strong relationship between openness to experience and acceptance of evolution in the previous cross-sectional survey study (Dunk et al., 2017). It is possible that the current student population differs in their relative importance of the factors related to evolution acceptance when compared to the previous student population; this is explored in the general linear models and discussed below. If the importance of openness differs, it could be manifest in a “ceiling effect” whereby individuals in the current study already have a level of openness that has maximal impact on acceptance of evolution, and no increase has a measurable further effect. Alternate explanations are the possibility that the change in openness to experience has a delayed effect on acceptance of evolution, or the possibility that openness to experience only has an effect for larger changes beyond those seen here.

We similarly found changes in intrinsic religiosity to have little relationship with changes in acceptance of evolution. Though the relationship was significant and in the expected direction (with decreasing intrinsic religiosity being associated with increasing acceptance of evolution), less than 4 percent of the variation in change in acceptance of evolution could be explained by changes in intrinsic religiosity. It is important to note this finding does not mean that intrinsic religiosity is not an important factor in acceptance of evolution (see general linear models), but rather that *changes in the level* of intrinsic religiosity do not relate strongly to changes in acceptance of evolution. These changes in evolutionary acceptance thus occur mostly independent of religiosity, which is counterintuitive compared to the strong importance of religiosity found in previous cross-sectional studies (Dunk et al., 2017; Glaze et al., 2015). This finding is consistent, however, with the possibility of students reducing their perceived conflict between evolution and religion throughout the semester (Barnes, Elser, & Brownell, 2017).

Finally, we found that increases in biological knowledge were moderately and significantly associated with increases in evolution acceptance. Specifically, two factors from the short form of the evolutionary attitudes and literacy survey (Short & Hawley, 2012), evolutionary knowledge and genetic literacy, had this strong positive relationship, while a third factor, evolutionary misconceptions, was not significantly related. It is somewhat surprising that observed changes in evolutionary misconceptions are not associated with changes in evolution acceptance. However, the instrument measures only a few, very specific misconceptions; it is possible the student population in the present study has other misconceptions that, if measured, would have a stronger relationship. Further, while we expected changes in both evolutionary knowledge and genetic literacy (as in Miller, Scott, & Okamoto, 2006) to be related to changes in evolution acceptance, we did not expect changes in genetic literacy (knowledge) to have a stronger, more significant, impact. While genetic mechanisms underlie so much evolutionary change, it is possible that the somewhat more indirect nature of knowledge of genetic mechanisms leads to a stronger relationship with acceptance of evolution when compared to evolutionary knowledge because there is reduced opportunity for backfire effects such as belief polarization (see Lewandowsky, Ecker, Seifert, Schwarz, & Cook, 2012 for summary).

### (iii) Pre-course and post-course general linear modeling

The differences between the models created from the pre-course and post-course survey administrations showed a number of important changes across the year. Looking at the summed effect size of each general group across the year, we see a marked decrease of the influence of political variables and a marked increase in the influence of nature of science understanding variables on acceptance of evolution when comparing the end-of-the-year model to the start-of-the-year model. We also see a decrease in the influence of knowledge and religious variables and in increase in the influence of demographic variables. However, beyond this broad view, it is useful to look in more detail at the changes in the effect size of individual model terms from the early fall to late spring models.

As mentioned above, the impact of religious variables went down over the year, but we expected a more profound change than that seen. Interestingly, while the overall impact of religious variables decreased from 9.7% to 7.2% of variance in acceptance of evolution explained, the individual variables shifted much more considerably. Specifically, religious affiliation, a very general coding of religious denomination, went from explaining over 8% of variance in early fall (the most of any single terms in the model) to being an insignificant model term in spring. In its stead, the number of religious friends an individual reported having (of any religion) went from being an insignificant variable in fall to explaining over 4.7% of the variance in spring. To us, this signals that these individuals may be shifting in their understanding of the interplay between religion and evolution throughout the year. That is, individuals start out the year with ideas about the relationship between evolution and religion that is guided mostly by their denomination; however, after a year of interaction with people of different denominations and faiths, it tends to be the case that a more religiously diverse community of friends guides their understanding, and that this impact is less strong than that seen by religious affiliation at the start of the year. The importance of religious friends after a year of biology may also mirror the recent finding that gains in acceptance of evolution are only significantly impacted by in-group identity (Walker, Wassenberg, Franta, & Cotner, 2017).

Interestingly, openness to experience did not have a strong enough relationship with acceptance of evolution to be included in either final model in this yearlong study, despite its strong relationship with acceptance of evolution in previously published models (Dunk et al., 2017; Hawley et al., 2011). One possibility is that there was significant overlap between the variance explained by openness to experience and the political variables in the full model, leaving no meaningful variance left for openness to explain after the political variables were included. This is consistent with findings that show openness to experience is highly correlated with political ideology (Van Hiel, Kossowska, & Mervielde, 2000). It is also possible, as discussed previously, that openness to experience is related to acceptance of evolution only in certain cases or at certain levels not present in our sample.

Though they decreased in importance from fall to spring, political variables in both models explained a large amount of the variation in acceptance of evolution in both the beginning of fall and the end of spring. While this may be unsurprising to many readers, we expected a lesser role for political variables compared to more nuanced psychological variables in the model. Additionally, previous research (Carter & Wiles, 2014) found that political identity was potent in explaining attitudes towards climate change, but had a smaller role in evolution acceptance. We are unsure if the difference between the previous study and the current one is due to a difference in the measures or model employed or a trend of increasing political polarization in acceptance of evolution, at least among students at the studied university.

When looking at the individual model terms for the political variables, we were surprised to find that two seemingly similar variables explained substantial, independent portions of variance in acceptance of evolution. We are unsure what substantive differences exist between identification as democrat, republican, or independent versus identification of general political views on a scale from conservative to liberal to drive this finding, but it exists and was robust enough to find at both the beginning and end of the year. Further research seeking to understand evolution acceptance should be sure to include both measures of political affiliation, so we can have comparison samples to begin to understand how these variables are affecting individuals’ acceptance of evolution.

These two political variables combined explained the greatest amount of variation in evolution acceptance of any grouping of variables in the early fall, with over 12% of variance explained. By late spring, this had decreased to 8.6% of variance explained: still a substantial portion, but an amount more equal to all general groupings. Intriguingly, the changes in the political variables between early fall and late spring were unequally divided between the two variables; general political views (again, conservatism vs liberalism) retained their impact throughout the year, while the impact of political party reduced from 7.4% to just over 4% of variance explained. While this general reduction seems to fit an interpretation of evolution education and/or increasing epistemological sophistication reducing a reliance on identities for understanding of scientific phenomena, we are unsure why this impact would solely be felt in party ID and not political views more broadly.

As a group, variables that indicate biological content knowledge did not shift appreciably in their impact on evolution acceptance from early fall to late spring, only decreasing by less than 1% of variation in evolution acceptance explained– the individual terms changed considerably more, though. By the end of the year, the impact of genetic literacy increased by more than two-fold compared to the beginning of the year, from 1.2% to 3.1% variance explained. This increase was coupled with a decrease in the impact of the number of biology classes taken in college, which changed from explaining almost 3% of variation in acceptance of evolution in early fall to no longer being a significant model term in spring. Evolutionary knowledge, on the other hand, explained about 4% of the variation in acceptance of evolution in both fall and spring models.

The shifts in knowledge variables show that a year of introductory biology instruction mitigates the impact that unequal prior college biology instruction had on evolution acceptance at the beginning of the fall semester. This, in turn, was replaced by an increased importance in genetic literacy, which may be due to an increased understanding amongst the more genetically literate of how evolutionary change can be documented by small-scale changes in population genetics (although this interpretation is speculative and needs to be explored in future research). The impact of genetic literacy on evolution acceptance has been recently found in a UK precollege population (Mead, Hejmadi, & Hurst, 2017), and was also found in an international, multifactorial study of evolution attitudes in the general public (Miller et al., 2006).

At neither time point did evolutionary misconceptions from the EALS-SF have a significant impact on an individuals’ acceptance of evolution when controlling for other variables. This is in line with the weak impact changes in evolutionary misconceptions had on changes in acceptance of evolution in this sample. It is possible that the instrument used did not include enough relevant misconceptions to accurately gauge the impact these misunderstandings of evolution have on evolution acceptance. However, we think it is also possible that measuring misconceptions is an ineffective way to gauge evolutionary acceptance in general, as students may accept evolution even while retaining misconceptions. Even biology instructors have been found to have a fairly high number of misconceptions about evolution (Nehm & Schonfeld, 2007), and such misconceptions can often be difficult to unseat (Nehm & Reilly, 2007).

Broadly, the impact of demographic variables on acceptance of evolution increased from early fall to late spring (from 5.4% to 7.1% variance explained). This trend was in the opposite direction of that expected, as we predicted that demographic variables would represent preparation and exposure to evolution, two things that a semester of introductory biology would tend to efface the effects of. One of the model terms, race/ethnicity, did behave this way. That is, the overall amount of variation in evolution acceptance explained by a respondent’s race/ethnicity decreased from over 5.3% in early fall to 3.5% in late spring. This trend is in a direction that is promising, but we are somewhat disappointed that the effect of race and ethnicity did not totally disappear (keeping in mind that differences we may expect to see between racial or ethnic groups, such as those due to differing religious affiliations, were already in the model). One possibility is that race and ethnicity in the current student population is associated with other socioeconomic variables that have a general negative effect on access to education; this is supported in theory by the finding that the year of instruction ended with a greater similarity in average MATE score between self-reported racial or ethnic identities. It could also explain why racial and ethnic identity was not significant in our previous study (Dunk et al., 2017), as that study used a student population that might be expected to be more equitable with respect to socioeconomic distribution between racial and ethnic identities.

We found an even more surprising change between fall and spring with respect to the other significant demographic model term, which measured childhood informal science exposure. This variable went from being insignificant in fall to explaining 3.5% of the variation in acceptance of evolution in the spring. We would have expected that a variable such as this would be more important in the fall as it seems to measure in some way students’ level of preparation. We are unsure why the results are in the opposite direction, but suggest that perhaps the increase is due to some change in an unmeasured variable. For example, perhaps individuals with more childhood science exposure were able to take better advantage of the instruction throughout the semester, and thus this exposure is not important so much in itself but in the way it allowed students to receive new information.

Finally, we look at the nature of science variables. As discussed above, as a group, nature of science variables’ effect size increased considerably from fall to spring (5.1% of variance explained in early fall compared to 8.8% explained in late spring). This increase led to an understanding of the nature of science to be the most important group of variables explaining variation in evolution acceptance in the spring (although all groups were within a fairly small percent of difference from each other). Looking at the individual model terms, there is notable difference between the two models. An understanding of the nature of science as unified was significant throughout the year, although it became much more impactful by the end of the year (increasing from 2.5 to 4.4 percent of variance explained). However, an understanding of science as amoral was only important in the early fall and was not included in the spring model (due to insignificance in the previous step’s “full model”). Likewise, an understanding of science as a process that is composed of, and requires, testable predictions was not eligible to be included in the model at the beginning of the year, but was very significant by the end of the year, explaining 4.4% of the variation in acceptance of evolution.

While we did not have specific predictions about how the importance of the individual components of the NSKS may change throughout the year, we think the results might fit well with a move from a naïve to a more mature understanding of the nature of science and evolutionary biology. That is, some individuals at the start of the year are influenced by their prior conceptions that science has a moral component and can make statements that compete in that arena. This would be especially problematic for religious students that use their religion as a moral guide if they feel that scientific knowledge is a replacement for this aspect of their faith; such a problem may lead to such students to feel uncomfortable in a biology classroom, which can lead to disengagement (Barnes, Truong, & Brownell, 2017). In contrast, an understanding of the testable nature of science leads to an understanding of the distinction between science and other forms of knowing– an understanding that scientific claims require testable hypotheses and that the majority of religious claims do not qualify as science due to this distinction. The testable nature of science (under the similar understanding of tentativeness) has often been associated with increased evolution acceptance (e.g.,Borgerding, Deniz, & Anderson, 2017).

In the past five years, researchers of evolution education have found that individual relationships exist between acceptance of evolution and the general groups of factors such as knowledge variables (Carter & Wiles, 2014; Cofré, Cuevas, & Becerra, 2017; Mead et al., 2017), political variables (Cotner et al., 2014), nature of science variables (Carter & Wiles, 2014; Cofré, Santibáñez, et al., 2017; Cofré, Cuevas, et al., 2017), and religious variables (Carter & Wiles, 2014), which are all general categories of variables we found significant in our analysis as well. In addition, many recent authors have found that psychological measures such as need for cognition (Kurdna, Shore, & Wassenberg, 2015) and epistemological types (Borgerding et al., 2017) to affect acceptance of evolution; we did not find a relation between acceptance of evolution and our psychological measure, openness to experience.

Comparing our study to multifactorial studies published within the past five years as well as another recent and well cited paper places our findings in even better context. When accounting for other variables, our study and others have found evolution acceptance to be significantly impacted by knowledge of evolution (Dorner, 2016; Dunk et al., 2017; Glaze et al., 2015; Mead et al., 2017; Weisberg, Landrum, Metz, & Weisberg, 2018), genetic knowledge (Mead et al., 2017; Miller et al., 2006), political variables (Miller et al., 2006; Walker et al., 2017; Weisberg et al., 2018), nature of science variables (Dorner, 2016; Dunk et al., 2017; Glaze et al., 2015), and religious variables (Dunk et al., 2017; Glaze et al., 2015; Miller et al., 2006; Rissler et al., 2014; Weisberg et al., 2018), as well as demographic variables such as race/ethnicity (Walker et al., 2017). However, differences exist as well. Others have found evolution acceptance to be impacted by age (Miller et al., 2006; Weisberg et al., 2018) and gender (Miller et al., 2006), but our model (as well as a previous one by us; Dunk et al., 2017) found no impact of either of these. Further, other studies find an impact of general educational attainment (Miller et al., 2006; Rissler et al., 2014; Walker et al., 2017; Weisberg et al., 2018), which we did not test directly; our closest proxy was number of college biology courses taken, which we found to be important in the beginning of the year, but not the end of the year.

## Limitations

While the findings in this paper are supported by robust statistical evidence, all studies are only as applicable as their study population. With that in mind, we acknowledge that these findings are from an undergraduate student population, which is further limited by a plurality of students being white and female. We further acknowledge the limitation of conducting the study at a private northeastern university; although many of our results are supported by previous work of ours at a public midwestern university, we encourage others to conduct similar studies in diverse academic settings and would be open to collaborations to do so. We also acknowledge the limitation of using only students in introductory biology. We are currently conducting a study that will explore similar questions using a more general student population. We would encourage others to do the same, as well as to explore the differences between novice and experienced biology students. The final limitation we would like to address is the notion of causality in our study. It should be noted that none of the relationships described above meet a strict notion of causality; we have shown important associations between variables, but the direction of that relationship is not tested. It is possible some causal language has made its way into our descriptions, and we apologize if that is the case; nonetheless, our results do show significant interactions between the variables discussed and acceptance of evolution. We feel that the results of significant correlations between change in acceptance of evolution and change in other variables sets a strong case for the potential that the associated variables do indeed cause a change; however, we acknowledge that further studies need to be done to establish directional causality, and we enthusiastically encourage such efforts.

## Conclusions

Despite the described changes in importance of variables throughout the semester, our data finds that all general groups of variables we defined (political, religious, nature of science, knowledge, and demographic) make a substantial contribution to explaining the variance in evolution acceptance. Further, these variables are similar to those found important in many of the studies of evolution education and acceptance conducted in the past five years in a variety of settings. Here, we have extended those studies by analyzing models both at the beginning and end of a year of biology instruction and showing how the impact of different factors on evolution acceptance change throughout the year. In addition, we have provided the beginnings of a causal link by showing how the change in associated variables, most significantly nature of science variables, is related to the change in acceptance of evolution.

We found a series of changes that occurred in the relationships between acceptance of evolution and associated variables between the beginning and end of a year of general biology instruction. Most notably, we found a sharp decrease in the importance of political associations on evolution acceptance, along with a decrease in religious variables. This was complimented by an increased importance in an understanding of the nature of science on the acceptance of evolution. Looking specifically at changes across the year, we found that changes in understanding the nature of science, genetic literacy, and evolutionary knowledge were strongly and significantly correlated with changes in evolution acceptance, indicating that these are all very fruitful potential targets for interventions designed to increase the acceptance of evolution.

We undertook this study to improve upon previous studies, but also to set a new baseline for further explorations of acceptance of evolution, especially in a longitudinal format. This baseline will allow further research of ours and others to explore the similarities and differences between different groups in acceptance of evolution (such as between students at different types of institutions, and ideally, between undergraduate students and different segments of the general population). Additionally, findings in this study have the potential to have direct applications to curriculum development and conceptual change research.

## Supplement

**Table S1.** Question wording of demographic variables.

|  |
| --- |
| 1. What is your gender identity? |
| *Free Response* |
| 2. What is your current age (in years)? |
| *Free Response* |
| 3. Do you consider yourself to be "Pre-med"? |
| *A. Yes B. No* |
| 4. What is your major or intended major? (NOTE: Pre-med is not a major) |
| *Free Response* |
| 5. Which of the following best describe you? Select all that apply. |
| *A. American Indian or Alaska Native B. Asian C. Black or African American D. Hispanic or Latino E. White F. Other* |
| 6. If you selected "Other" in the previous question please state your race/ethnicity in the text box here. |
| *Free Response* |
| 7. If you are from the United States, please type the state or territory you are from. If you are not from the United States, please type the country you are from. |
| *Free Response* |
| 8. Which term best describes where you grew up? |
| *A. Urban B. Suburban C. Rural* |
| 9. Growing up, how often were you exposed to science outside of school (e.g., by visiting museums, science centers, etc.)? |
| *A. Almost Never B. Rarely C. Somewhat Rarely D. Somewhat Often E. Very Often* |
| 10. How would you rank your interest in science in general? |
| *A. Not at all interested B. Mostly Uninterested C. Neutral D. Somewhat interested E. Very Interested* |
| 11. How many science classes have you taken in college (excluding this one)? |
| *Free Response* |
| 12. How many biology classes have you taken in college (excluding this one)? |
| *Free Response* |
| 13. What is your mother's highest level of education? |
| *A. Never attended school or only attended kindergarten B. Grades 1 through 8 (Elementary) C. Grades 9 through 11 (Some high school) D. Grade 12 or GED (High school graduate)* |
| *E. Attended college but did not graduate*<br>*F. Associate's or technical degree*<br>*G. College graduate (Bachelor's degree)*<br>*H. Post-bachelor's degree (Graduate school, Law school, Medical school)* |
| 14. What is your father's highest level of education? |
| *A. Never attended school or only attended kindergarten*<br>*B. Grades 1 through 8 (Elementary)*<br>*C. Grades 9 through 11 (Some high school)*<br>*D. Grade 12 or GED (High school graduate)*<br>*E. Attended college but did not graduate*<br>*F. Associate's or technical degree*<br>*G. College graduate (Bachelor's degree)*<br>*H. Post-bachelor's degree (Graduate school, Law school, Medical school)* |
| 15. What, if any, is your religious affiliation? |
| *A. Agnostic B. Atheist C. Buddhist D. Catholic E. Evangelical Christian*<br>*F. Hindu G. Jewish H. Mainline Protestant I. Muslim*<br>*J. Non-denominational Christian K. Spiritual but not religious L. Other* |
| 16. If you answered "Other" in the previous question, please use this text box to type in your religious denomination. You may also use this space to clarify or add detail to your response regardless of your choice above. |
| *Free Response* |
| 17. How active do you consider yourself to be in the practice of your religious preference? |
| *A. Not active B. Not very active C. Somewhat active*<br>*D. Very active E. Does not apply* |
| 18. In general, how would you describe your political views? |
| *A. Strongly liberal B. Somewhat liberal C. Moderate/ Middle of the road*<br>*D. Somewhat conservative E. Strongly conservative* |
| 19. Politically, what are your views on most social issues (e.g., immigration, capital punishment, or marriage equality): |
| *A. Strongly liberal B. Somewhat liberal C. Moderate/ Middle of the road*<br>*D. Somewhat conservative E. Strongly conservative* |
| 20. Politically, what are your views on most fiscal issues (e.g., government spending, trade regulation, or economic regulation): |
| *A. Strongly liberal B. Somewhat liberal C. Moderate/ Middle of the road*<br>*D. Somewhat conservative E. Strongly conservative* |
| 21. Generally speaking, do you consider yourself to be a(n): |
| *A. Strong Democrat B. Not-so-strong Democrat*<br>*C. Independent-leaning Democrat D. Independent*<br>*E. Independent-leaning Republican F. Not-so-strong Republican*<br>*G. Strong Republican H. Other (see next question) I. Don't Know* |
| 22. If you answered "Other" to the previous question please use the text box here to type your political party affiliation. If you made a selection in the previous question please leave this blank. |
| *Free Response* |

**Table S2.** Frequency tables for categorical variables not presented in the main text. Data is from fall survey administration.*

| Variable | Category | Number<sup>1</sup> | Percent |
| --- | --- | --- | --- |
| Pre-Med Student |  |  |  |
|  | Yes | 181 | 34.4 |
|  | No | 345 | 65.6 |
| Census Region |  |  |  |
|  | International | 35 | 6.9 |
|  | Midwest | 21 | 4.1 |
|  | Northeast | 351 | 69.2 |
|  | South | 44 | 8.7 |
|  | West | 44 | 8.7 |
|  | Puerto Rico | 4 | 0.8 |
|  | Other | 8 | 1.6 |
| Science Interest |  |  |  |
|  | Not at all interested | 7 | 1.3 |
|  | Mostly uninterested | 33 | 6.3 |
|  | Neutral | 70 | 13.4 |
|  | Somewhat interested | 165 | 31.7 |
|  | Very interested | 246 | 47.2 |
| Mother's Education Level |  |  |  |
|  | Never attended school or only attended kindergarten | 0 | 0.0 |
|  | Grades 1 through 8 (Elementary) | 5 | 1.0 |
|  | Grades 9 through 11 (Some high school) | 12 | 2.3 |
|  | Grade 12 or GED (High school graduate) | 85 | 16.3 |
|  | Attended college but did not graduate | 38 | 7.3 |
|  | Associate's or technical degree | 49 | 9.4 |
|  | College graduate (Bachelor's degree) | 174 | 33.3 |
|  | Post-bachelor's degree (Graduate school, law school, medical school) | 154 | 29.4 |
|  | Does not apply | 6 | 1.1 |
| Father's Education Level |  |  |  |
|  | Never attended school or only attended kindergarten | 1 | 0.2 |
|  | Grades 1 through 8 (Elementary) | 9 | 1.7 |
|  | Grades 9 through 11 (Some high school) | 15 | 2.9 |
|  | Grade 12 or GED (High school graduate) | 88 | 16.8 |
|  | Attended college but did not graduate | 36 | 6.9 |
|  | Associate's or technical degree | 45 | 8.6 |
|  | College graduate (Bachelor's degree) | 146 | 27.9 |
|  | Post-bachelor's degree (Graduate school, law school, medical school) | 162 | 31.0 |
|  | Does not apply | 21 | 4.0 |
| Social Political Views |  |  |  |
|  | Strongly Conservative | 12 | 2.3 |
|  | Somewhat Conservative | 45 | 8.7 |
|  | Moderate/ Middle of the Road | 162 | 31.2 |
|  | Somewhat Liberal | 161 | 31.0 |
|  | Strongly Liberal | 140 | 26.9 |
| Fiscal Political Views |  |  |  |
|  | Strongly Conservative | 31 | 6.0 |
|  | Somewhat Conservative | 98 | 18.8 |
|  | Moderate/ Middle of the Road | 241 | 46.3 |
|  | Somewhat Liberal | 112 | 21.5 |
|  | Strongly Liberal | 39 | 7.5 |
| Number of Similarly Religious Friends |  |  |  |
|  | 0 | 52 | 10.2 |
|  | 1 | 58 | 11.3 |
|  | 2 | 102 | 19.9 |
|  | 3 | 122 | 23.8 |
|  | 4 | 92 | 18.0 |
|  | 5 | 86 | 16.8 |

\*See table 3 in main text for the remaining categorical variables.
<sup>1</sup>Numbers in each category may not add to the same total due to nonresponse. Nonresponses are not included.

**Table S3.**
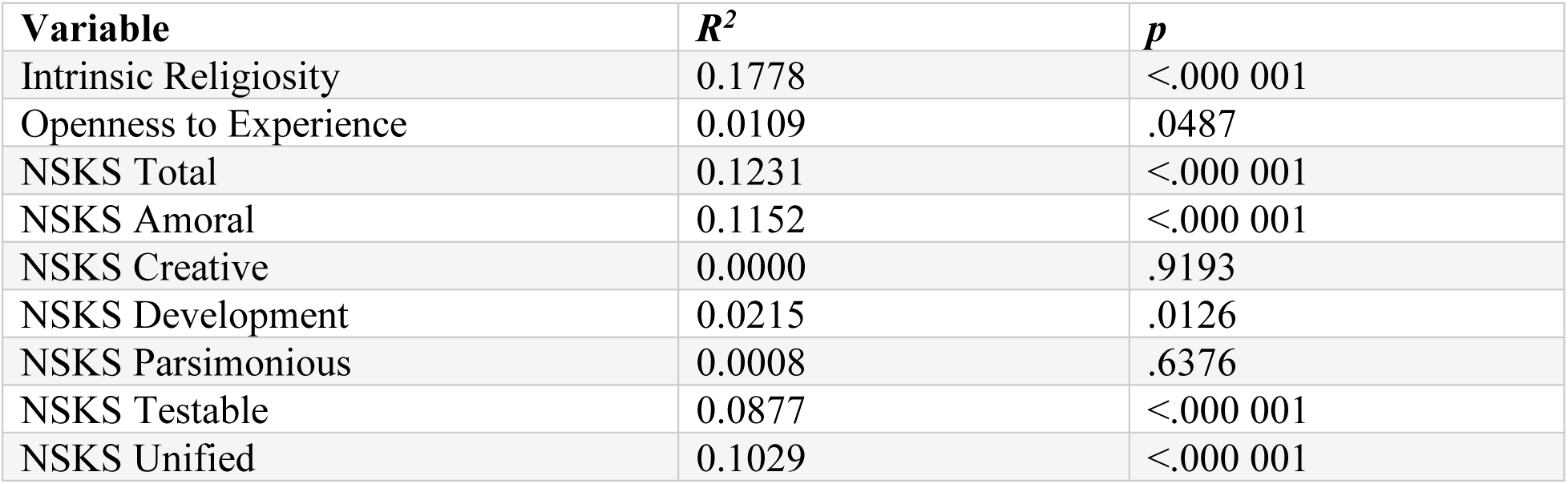
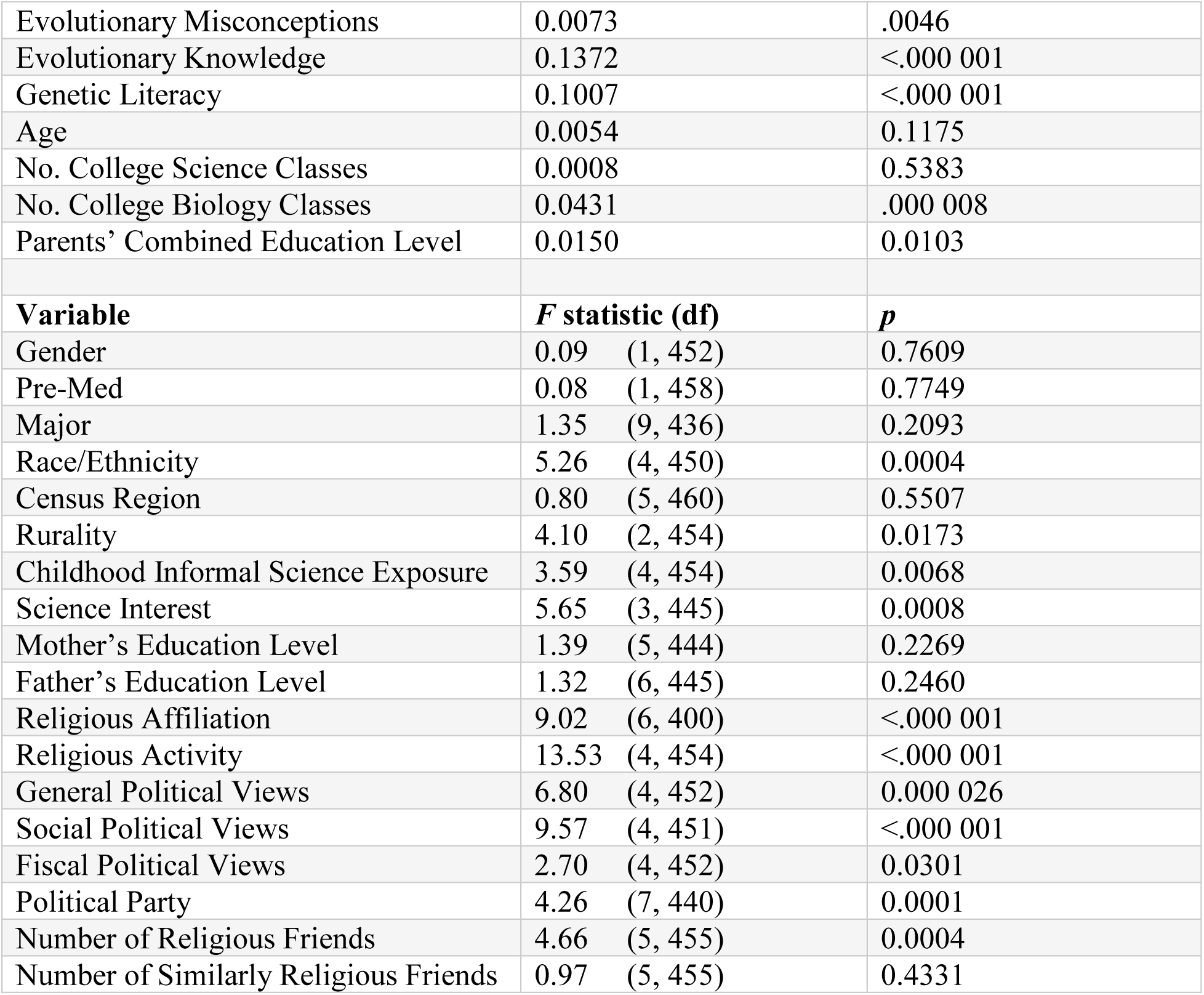
Results of individual correlations or ANOVAs of given variables on MATE score in fall semester.

| Variable | $R^2$ | $p$ |
| --- | --- | --- |
| Intrinsic Religiosity | 0.1778 | <.000 001 |
| Openness to Experience | 0.0109 | .0487 |
| NSKS Total | 0.1231 | <.000 001 |
| NSKS Amoral | 0.1152 | <.000 001 |
| NSKS Creative | 0.0000 | .9193 |
| NSKS Development | 0.0215 | .0126 |
| NSKS Parsimonious | 0.0008 | .6376 |
| NSKS Testable | 0.0877 | <.000 001 |
| NSKS Unified | 0.1029 | <.000 001 |
| Evolutionary Misconceptions | 0.0073 | .0046 |
| Evolutionary Knowledge | 0.1372 | <.000 001 |
| Genetic Literacy | 0.1007 | <.000 001 |
| Age | 0.0054 | 0.1175 |
| No. College Science Classes | 0.0008 | 0.5383 |
| No. College Biology Classes | 0.0431 | .000 008 |
| Parents' Combined Education Level | 0.0150 | 0.0103 |
| **Variable** | **F statistic (df)** | **p** |
| Gender | 0.09 (1, 452) | 0.7609 |
| Pre-Med | 0.08 (1, 458) | 0.7749 |
| Major | 1.35 (9, 436) | 0.2093 |
| Race/Ethnicity | 5.26 (4, 450) | 0.0004 |
| Census Region | 0.80 (5, 460) | 0.5507 |
| Rurality | 4.10 (2, 454) | 0.0173 |
| Childhood Informal Science Exposure | 3.59 (4, 454) | 0.0068 |
| Science Interest | 5.65 (3, 445) | 0.0008 |
| Mother's Education Level | 1.39 (5, 444) | 0.2269 |
| Father's Education Level | 1.32 (6, 445) | 0.2460 |
| Religious Affiliation | 9.02 (6, 400) | <.000 001 |
| Religious Activity | 13.53 (4, 454) | <.000 001 |
| General Political Views | 6.80 (4, 452) | 0.000 026 |
| Social Political Views | 9.57 (4, 451) | <.000 001 |
| Fiscal Political Views | 2.70 (4, 452) | 0.0301 |
| Political Party | 4.26 (7, 440) | 0.0001 |
| Number of Religious Friends | 4.66 (5, 455) | 0.0004 |
| Number of Similarly Religious Friends | 0.97 (5, 455) | 0.4331 |

**Table S4.**
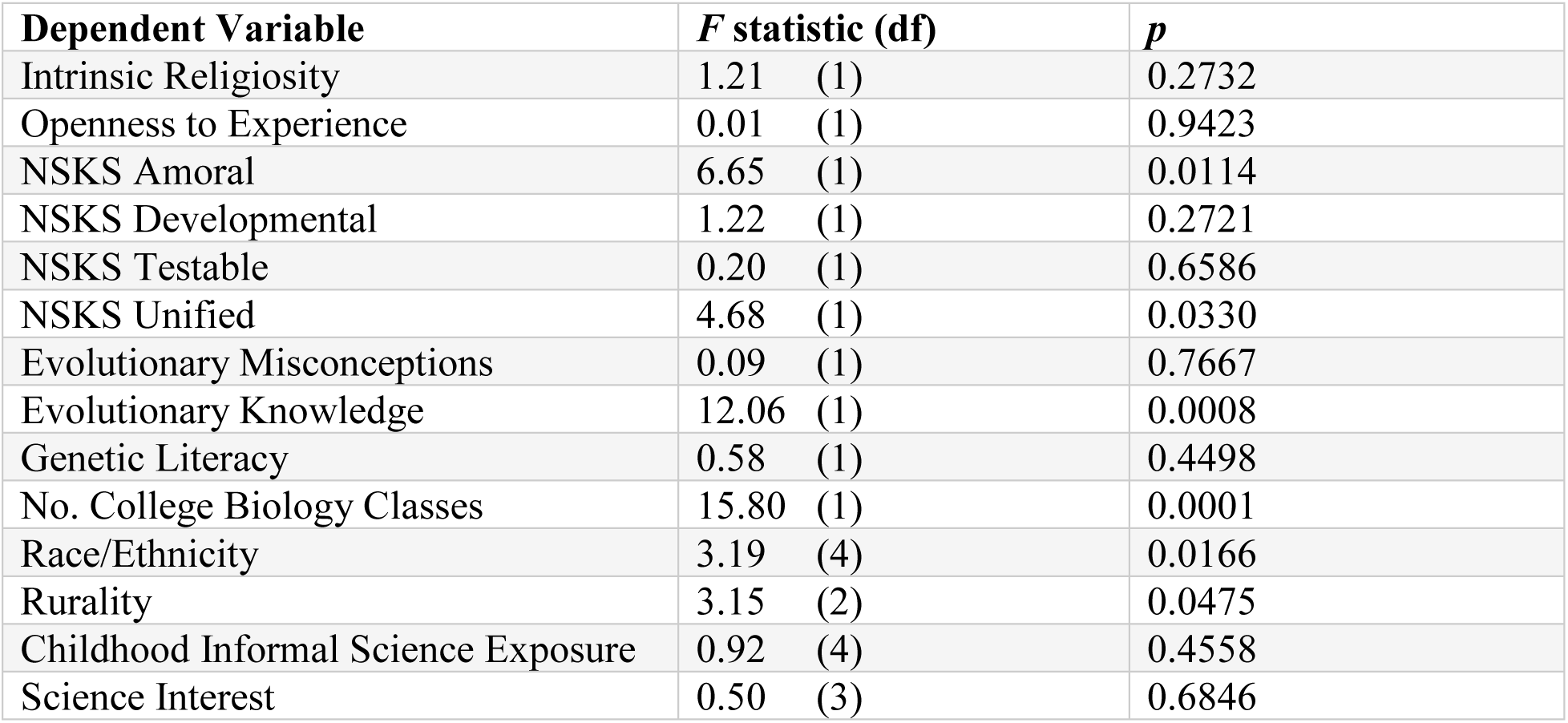
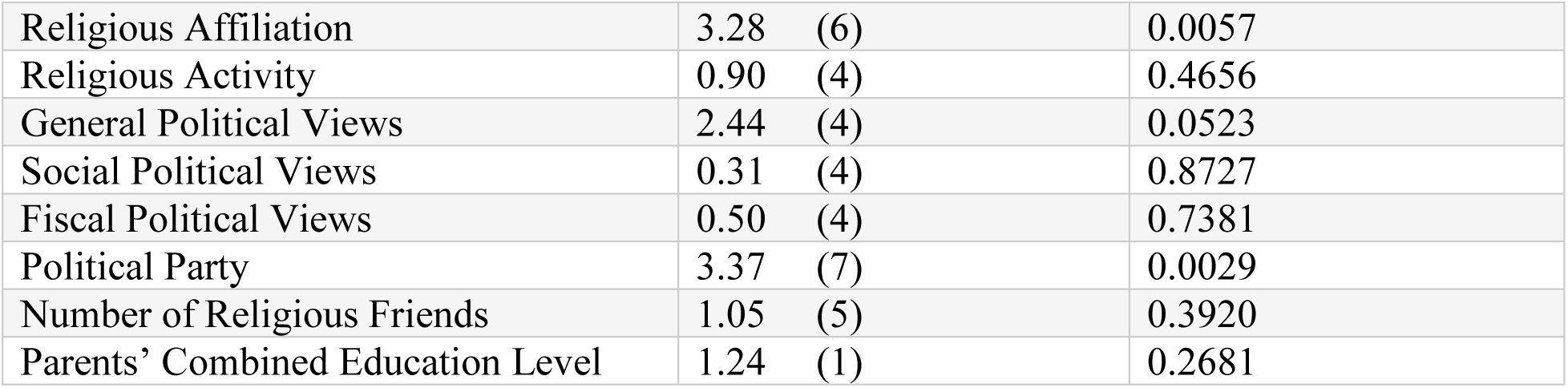
Results of “full model” GLM of given variables on MATE score in fall semester.

|  |  |  |
| --- | --- | --- |
| **Dependent Variable** | **F statistic (df)** | **p** |
| Intrinsic Religiosity | 1.21 (1) | 0.2732 |
| Openness to Experience | 0.01 (1) | 0.9423 |
| NSKS Amoral | 6.65 (1) | 0.0114 |
| NSKS Developmental | 1.22 (1) | 0.2721 |
| NSKS Testable | 0.20 (1) | 0.6586 |
| NSKS Unified | 4.68 (1) | 0.0330 |
| Evolutionary Misconceptions | 0.09 (1) | 0.7667 |
| Evolutionary Knowledge | 12.06 (1) | 0.0008 |
| Genetic Literacy | 0.58 (1) | 0.4498 |
| No. College Biology Classes | 15.80 (1) | 0.0001 |
| Race/Ethnicity | 3.19 (4) | 0.0166 |
| Rurality | 3.15 (2) | 0.0475 |
| Childhood Informal Science Exposure | 0.92 (4) | 0.4558 |
| Science Interest | 0.50 (3) | 0.6846 |
| Religious Affiliation | 3.28 (6) | 0.0057 |
| Religious Activity | 0.90 (4) | 0.4656 |
| General Political Views | 2.44 (4) | 0.0523 |
| Social Political Views | 0.31 (4) | 0.8727 |
| Fiscal Political Views | 0.50 (4) | 0.7381 |
| Political Party | 3.37 (7) | 0.0029 |
| Number of Religious Friends | 1.05 (5) | 0.3920 |
| Parents' Combined Education Level | 1.24 (1) | 0.2681 |

**Table S5.**
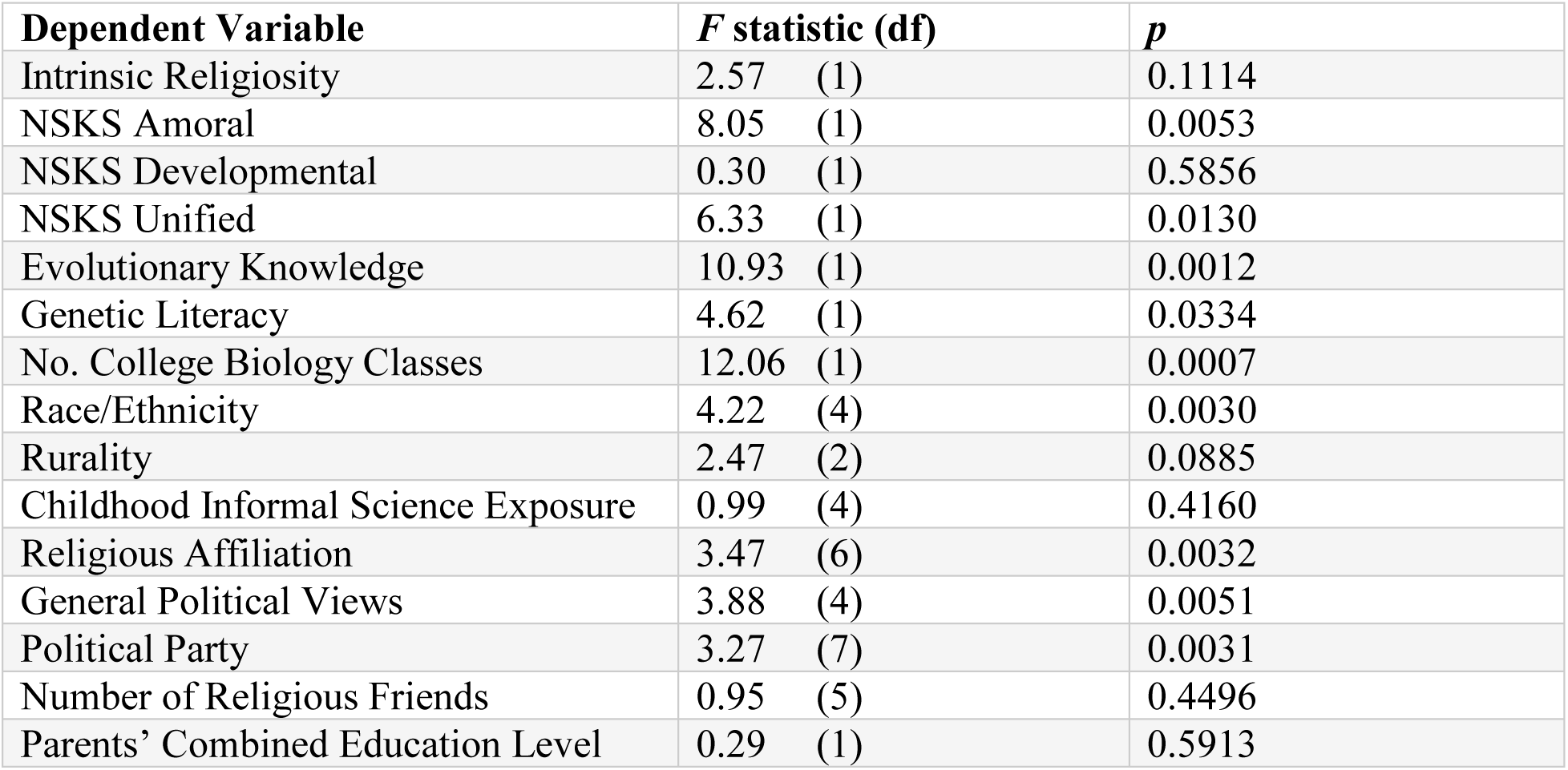
Results of “intermediate model” GLM of given variables on MATE score in fall semester.

| **Dependent Variable** | **F statistic (df)** | **p** |
| --- | --- | --- |
| Intrinsic Religiosity | 2.57 (1) | 0.1114 |
| NSKS Amoral | 8.05 (1) | 0.0053 |
| NSKS Developmental | 0.30 (1) | 0.5856 |
| NSKS Unified | 6.33 (1) | 0.0130 |
| Evolutionary Knowledge | 10.93 (1) | 0.0012 |
| Genetic Literacy | 4.62 (1) | 0.0334 |
| No. College Biology Classes | 12.06 (1) | 0.0007 |
| Race/Ethnicity | 4.22 (4) | 0.0030 |
| Rurality | 2.47 (2) | 0.0885 |
| Childhood Informal Science Exposure | 0.99 (4) | 0.4160 |
| Religious Affiliation | 3.47 (6) | 0.0032 |
| General Political Views | 3.88 (4) | 0.0051 |
| Political Party | 3.27 (7) | 0.0031 |
| Number of Religious Friends | 0.95 (5) | 0.4496 |
| Parents' Combined Education Level | 0.29 (1) | 0.5913 |

**Table S6.** Results of individual correlations or ANOVAs of given variables on MATE score in spring semester.

| **Variable** | **R<sup>2</sup>** | **p** |
| --- | --- | --- |
| Intrinsic Religiosity | 0.1668 | <.000 001 |
| Openness to Experience | 0.0459 | 0.0002 |
| NSKS Total | 0.4096 | <.000 001 |
| NSKS Amoral | 0.1537 | <.000 001 |
| NSKS Creative | 0.0262 | 0.0062 |
| NSKS Development | 0.2942 | <.000 001 |
| NSKS Parsimonious | 0.0321 | 0.0024 |
| NSKS Testable | 0.3293 | <.000 001 |
| NSKS Unified | 0.4216 | <.000 001 |
| Evolutionary Misconceptions | 0.0539 | 0.000 024 |
| Evolutionary Knowledge | 0.3939 | <.000 001 |
| Genetic Literacy | 0.3702 | <.000 001 |
| Age | 0.0024 | 0.3844 |
| No. College Science Classes | 0.0003 | 0.7694 |
| No. College Biology Classes | 0.0048 | 0.2231 |
| Parents' Combined Education Level | 0.0025 | 0.3903 |
| **Variable** | **F statistic (df)** | **p** |
| Gender | 0.20 (1, 309) | 0.6539 |
| Pre Med | 0.62 (1, 311) | 0.4308 |
| Major | 0.98 (7, 289) | 0.4439 |
| Race/Ethnicity | 5.77 (4, 300) | 0.0002 |
| Region | 1.58 (4, 290) | 0.1791 |
| Rurality | 5.33 (2, 309) | 0.0053 |
| Childhood Informal Science Exposure | 3.90 (4, 306) | 0.0042 |
| Science Interest | 5.52 (4, 307) | 0.0003 |
| Mother's Education Level | 0.62 (4, 293) | 0.6472 |
| Father's Education Level | 0.79 (6, 302) | 0.5767 |
| Religious Affiliation | 5.11 (7, 275) | 0.000 018 |
| Religious Activity | 5.36 (4, 307) | 0.0004 |
| Political General | 4.79 (4, 306) | 0.0009 |
| Political Social | 6.86 (4, 305) | 0.000 027 |
| Political Fiscal | 2.42 (4, 304) | 0.0488 |
| Political Party | 3.48 (7, 292) | 0.0013 |
| Number of Religious Friends | 5.75 (5, 306) | 0.000 044 |
| Number of Similarly Religious Friends | 0.63 (5, 306) | 0.6782 |

**Table S7.** Results of “full model” GLM of given variables on MATE score in spring semester.

|  |  |  |
| --- | --- | --- |
| **Dependent Variable** | **F statistic (df)** | **p** |
| Intrinsic Religiosity | 10.87 (1) | 0.0012 |
| Openness to Experience | 0.18 (1) | 0.6693 |
| NSKS Amoral | 0.05 (1) | 0.8197 |
| NSKS Creative | 0.51 (1) | 0.4757 |
| NSKS Developmental | 0.11 (1) | 0.7387 |
| NSKS Parsimonious | 0.88 (1) | 0.3485 |
| NSKS Testable | 12.98 (1) | 0.0004 |
| NSKS Unified | 12.49 (1) | 0.0006 |
| Evolutionary Misconceptions | 1.37 (1) | 0.2444 |
| Evolutionary Knowledge | 17.29 (1) | 0.000 056 |
| Genetic Literacy | 5.98 (1) | 0.0157 |
| Race/Ethnicity | 3.78 (4) | 0.0060 |
| Rurality | 0.01 (2) | 0.9855 |
| Childhood Informal Science Exposure | 2.53 (4) | 0.0431 |
| Science Interest | 0.50 (4) | 0.7372 |
| Religious Affiliation | 1.63 (7) | 0.1316 |
| Religious Activity | 1.10 (4) | 0.3602 |
| Political General | 3.65 (4) | 0.0074 |
| Political Social | 0.22 (4) | 0.9292 |
| Political Fiscal | 0.64 (4) | 0.6354 |
| Political Party | 2.39 (7) | 0.0242 |
| Number of Religious Friends | 3.24 (5) | 0.0085 |

**Table S8.** Results of “intermediate model” GLM of given variables on MATE score in spring semester.

| **Dependent Variable** | ***F* statistic (df)** | ***p*** |
| --- | --- | --- |
| Intrinsic Religiosity | 8.68 (1) | 0.0037 |
| NSKS Creative | 0.24 (1) | 0.6218 |
| NSKS Parsimonious | 0.64 (1) | 0.4244 |
| NSKS Testable | 15.53 (1) | 0.0001 |
| NSKS Unified | 16.32 (1) | 0.000 080 |
| Evolutionary Misconceptions | 0.72 (1) | 0.3957 |
| Evolutionary Knowledge | 15.09 (1) | 0.0001 |
| Genetic Literacy | 11.55 (1) | 0.0008 |
| Race/Ethnicity | 3.30 (4) | 0.0123 |
| Childhood Informal Science Exposure | 3.40 (4) | 0.0105 |
| Religious Affiliation | 1.30 (7) | 0.2552 |
| Religious Activity | 1.08 (4) | 0.3665 |
| General Political Views | 4.32 (4) | 0.0023 |
| Political Party | 2.24 (7) | 0.0334 |
| Number of Religious Friends | 2.99 (5) | 0.0123 |

## References

Abdi, H. (2010). Holm’s sequential Bonferroni procedure. In N. Salkind (Ed.), Encyclopedia of research design (Vol. 2, pp. 573–577). Thousand Oaks, CA: SAGE Publications, Inc.

Akyol, G., Tekkaya, C., Sungur, S., & Traynor, A. (2012). Modeling the Interrelationships Among Pre-service Science Teachers’ Understanding and Acceptance of Evolution, Their Views on Nature of Science and Self-Efficacy Beliefs Regarding Teaching Evolution. Journal of Science Teacher Education, 23(8), 937–957. https://doi.org/10.1007/s10972-012-9296-x

Alters, B. J., & Alters, S. M. (2001). Defending Evolution in the Classroom: A Guide to the Creation/Evolution Controversy. Sudbury, MA: Jones and Bartlett Publishers.

Baker, J. O. (2013). Acceptance of evolution and support for teaching creationism in public schools: The conditional impact of educational attainment. Journal for the Scientific Study of Religion, 52(1), 216–228.

Barnes, M. E., & Brownell, S. E. (2016). Practices and Perspectives of College Instructors on Addressing Religious Beliefs When Teaching Evolution. Cell Biology Education, 15(2), ar18–ar18. https://doi.org/10.1187/cbe.15-11-0243

Barnes, M. E., Elser, J., & Brownell, S. E. (2017). Impact of a Short Evolution Module on Students’ Perceived Conflict between Religion and Evolution. The American Biology Teacher, 79(2), 104–111.

Barnes, M. E., Truong, J. M., & Brownell, S. E. (2017). Experiences of Judeo-Christian Students in Undergraduate Biology. Cell Biology Education, 16(1), ar15. https://doi.org/10.1187/cbe.16-04-0153

Borgerding, L. A., Deniz, H., & Anderson, E. S. (2017). Evolution acceptance and epistemological beliefs of college biology students. Journal of Research in Science Teaching, 54(4), 493–519. https://doi.org/10.1002/tea.21374

Brown, J. (2015). Measuring the acceptance of evolutionary theory: A profile of science majors in Texas 2-year colleges (Ph.D.). Texas A&M University, Commerce, TX.

Carter, B. E., & Wiles, J. R. (2014). Scientific consensus and social controversy: Exploring relationships between students’ conceptions of the nature of science, biological evolution, and global climate change. Evolution: Education and Outreach, 7, 6.

Cavallo, A. M., & McCall, D. (2008). Seeing may not mean believing: examining students’ understandings & beliefs in evolution. The American Biology Teacher, 70(9), 522–530.

Chan, K.-S. (2005). Exploring the dynamic interplay of college students’ conceptions of the nature of science. Asia-Pacific Forum on Science Learning and Teaching, 6(2), 1–16.

Cofré, H. L., Cuevas, E., & Becerra, B. (2017). The relationship between biology teachers’ understanding of the nature of science and the understanding and acceptance of the theory of evolution. International Journal of Science Education, 39(16), 2243–2260. https://doi.org/10.1080/09500693.2017.1373410

Cofré, H. L., Santibáñez, D. P., Jiménez, J. P., Spotorno, A., Carmona, F., Navarrete, K., & Vergara, C. A. (2017). The effect of teaching the nature of science on students’ acceptance and understanding of evolution: myth or reality? Journal of Biological Education, 1–14. https://doi.org/10.1080/00219266.2017.1326968

Cotner, S. H., Brooks, D. C., & Moore, R. (2014). Science and Society: Evolution and Student Voting Patterns. Reports of the National Center for Science Education, 34(6), 1–11.

Coyne, J. A. (2009). Why Evolution Is True. New York: Viking.

Deniz, H., Donnelly, L. A., & Yilmaz, I. (2008). Exploring the factors related to acceptance of evolutionary theory among Turkish preservice biology teachers: Toward a more informative conceptual ecology for biological evolution. Journal of Research in Science Teaching, 45(4), 420–443. https://doi.org/10.1002/tea.20223

Dobzhansky, T. (1973). Nothing in Biology Makes Sense except in the Light of Evolution. The American Biology Teacher, 35(3), 125–129. https://doi.org/10.2307/4444260

Dorner, M. A. (2016). Academic Factors that Predict Community College Students’ Acceptance of Evolution (Ph.D.). Chapman University, Orange, CA.

Dorner, M. A., & Scott, E. C. (2016). An exploration of instructor perceptions of community college students’ attitudes towards evolution. Evolution: Education and Outreach, 9(1). https://doi.org/10.1186/s12052-016-0055-x

Dunk, R. D. P., Petto, A. J., Wiles, J. R., & Campbell, B. C. (2017). A Multifactorial Analysis of Acceptance of Evolution. Evolution: Education and Outreach, 10, 4. https://doi.org/10.1186/s12052-017-0068-0

Eldredge, N. (2000). The Triumph of Evolution and the Failure of Creationism. New York: W. H. Freeman and Company.

Flower, P. (2006). Knowledge of and Attitudes toward Evolution in a Population of Community College Students. Forum on Public Policy Online, 2006(1), 1–12.

Folmer, V., Barbosa, N. de V., Soares, F. A., & Rocha, J. B. T. (2009). Experimental activities based on ill-structured problems improve Brazilian school students’ understanding of the nature of scientific knowledge. Revista Electrónica de Enseñanza de Las Ciencias, 8(1), 232–254.

Fox, J., & Monette, G. (1992). Generalized Collinearity Diagnostics. Journal of the American Statistical Association, 87(417), 178. https://doi.org/10.2307/2290467

Gallup. (2014, May 8). In U.S., 42% Believe Creationist View of Human Origins. Retrieved January 24, 2018, from http://news.gallup.com/poll/170822/believe-creationist-view-human-origins.aspx

Glaze, A. L., Goldston, M. J., & Dantzler, J. (2015). Evolution in the southeastern USA: Factors influencing acceptance and rejection in pre-service science teachers. International Journal of Science and Mathematics Education, 13(6), 1189–1209.

Graffin, G. (2003). Monism, atheism, and the naturalist world-view: Perspectives from evolutionary biology (Ph.D.). Cornell University, Ithaca, NY.

Grose, E. C., & Simpson, R. D. (1982). Attitudes of introductory college biology students towards evolution. Journal of Research in Science Teaching, 19(1), 15–24.

Grossman, W. E., & Fleet, C. M. (2017). Changes in acceptance of evolution in a college-level general education course. Journal of Biological Education, 51(4), 328–335. https://doi.org/10.1080/00219266.2016.1233128

Ha, M., Cha, H., & Ku, S. (2012). A comparative study of Korean and United States college students’ degree of religiosity, evolutionary interest, understanding and acceptance and their structures. Journal of The Korean Association For Science Education, 32(10), 1537–1550.

Hawley, P. H., Short, S. D., McCune, L. A., Osman, M. R., & Little, T. D. (2011). What’s the Matter with Kansas?: The Development and Confirmation of the Evolutionary Attitudes and Literacy Survey (EALS). Evolution: Education and Outreach, 4(1), 117–132. https://doi.org/10.1007/s12052-010-0294-1

Heddy, B. C., & Nadelson, L. S. (2013). The variables related to public acceptance of evolution in the United States. Evolution: Education and Outreach, 6(1), 1–14.

Hill, J. P. (2014). Rejecting evolution: The role of religion, education, and social networks. Journal for the Scientific Study of Religion, 53(3), 575–594.

Hill, P. C., & Hood, R. W. (Eds.). (1999). Measures of Religiosity. Birmingham, AL: Religious Education Press.

Howard, T. C., & Navarro, O. (2016). Critical race theory 20 years later: Where do we go from here? Urban Education, 51(3), 253–273.

Huitema, B. E. (2011). The analysis of covariance and alternatives (2nd ed.). Hoboken, NJ: John Wiley & Sons, Inc.

John, O. P., Naumann, L. P., & Soto, C. J. (2008). Paradigm Shift to the Integrative Big-Five Trait Taxonomy: History, Measurement, and Conceptual issues. In O. P. John, R. W. Robins, & L. A. Pervin (Eds.), Handbook of Personality Theory and Research (3rd ed., pp. 114–158). New York: Guilford Press.

Johnson, R. L., & Peeples, E. E. (1987). The Role of Scientific Understanding in College: Student Acceptance of Evolution. The American Biology Teacher, 49(2), 93–98. https://doi.org/10.2307/4448445

Kilic, K., Sungur, S., Cakiroglu, J., & Tekkaya, C. (2005). Ninth grade students’ understanding of the nature of scientific knowledge. Hacettepe Üniversitesi Eğitim Fakültesi Dergisi, 28(28).

Kurdna, J., Shore, M., & Wassenberg, D. (2015). Considering the Role of “Need for Cognition” in Students’ Acceptance of Climate Change & Evolution. The American Biology Teacher, 77(4), 250–257. https://doi.org/10.1525/abt.2015.77.4.4

Ladson-Billinngs, G., & Tate, W. (1995). Toward a critical race theory of education. Teachers College Record, 97(1), 47–68.

Lederman, N. G., Abd-El-Khalick, F., Bell, R. L., & Schwartz, R. S. (2002). Views of nature of science questionnaire: Toward valid and meaningful assessment of learners’ conceptions of nature of science. Journal of Research in Science Teaching, 39(6), 497–521. https://doi.org/10.1002/tea.10034

Lederman, N. G., Wade, P. D., & Bell, R. L. (1998). Assessing the nature of science: What is the nature of our assessments? Science and Education, 7, 595–615.

Lewandowsky, S., Ecker, U. K. H., Seifert, C. M., Schwarz, N., & Cook, J. (2012). Misinformation and Its Correction: Continued Influence and Successful Debiasing. Psychological Science in the Public Interest, 13(3), 106–131. https://doi.org/10.1177/1529100612451018

Lombrozo, T., Thanukos, A., & Weisberg, M. (2008). The Importance of Understanding the Nature of Science for Accepting Evolution. Evolution: Education and Outreach, 1(3), 290–298. https://doi.org/10.1007/s12052-008-0061-8

Lord, T., & Marino, S. (1993). How university students view the theory of evolution. Journal of College Science Teaching, 22(6), 353–357.

Lynn, C. D., Glaze, A. L., Evans, W. A., & Reed, L. K. (Eds.). (2017). Evolution Education in the American South: Culture, Politics, and Resources in and around Alabama. New York: Palgrave Macmillan.

Manwaring, K. F., Jensen, J. L., Gill, R. A., & Bybee, S. M. (2015). Influencing highly religious undergraduate perceptions of evolution: Mormons as a case study. Evolution: Education and Outreach, 8(1). https://doi.org/10.1186/s12052-015-0051-6

Marx, J. D., & Cummings, K. (2007). Normalized change. American Journal of Physics, 75(1), 87–91. https://doi.org/10.1119/1.2372468

Matthews, M. (1997). Editorial. Science & Education, 6(4), 323–329.

Mayr, E. (2001). What Evolution Is. New York: Basic Books.

Mazur, A. (2004). Believers and disbelievers in evolution. Politics and the Life Sciences, 23(2), 55–61.

Mead, R., Hejmadi, M., & Hurst, L. D. (2017). Teaching genetics prior to teaching evolution improves evolution understanding but not acceptance. PLOS Biology, 15(5), e2002255. https://doi.org/10.1371/journal.pbio.2002255

Meadows, L., Doster, E., & Jackson, D. F. (2000). Managing the Conflict between Evolution & Religion. The American Biology Teacher, 62(2), 102–107. https://doi.org/10.2307/4450848

Miller, J. D., Scott, E. C., & Okamoto, S. (2006). Public acceptance of evolution. Science, 313(5788), 765.

Moore, R., Brooks, D. C., & Cotner, S. (2011). The Relation of High School Biology Courses & Students’ Religious Beliefs to College Students’ Knowledge of Evolution. The American Biology Teacher, 73(4), 222–226. https://doi.org/10.1525/abt.2011.73.4.7

Nadelson, L. S., & Hardy, K. K. (2015). Trust in science and scientists and the acceptance of evolution. Evolution: Education and Outreach, 8(1). https://doi.org/10.1186/s12052-015-0037-4

Nadelson, L. S., & Southerland, S. (2012). A More Fine-Grained Measure of Students’ Acceptance of Evolution: Development of the Inventory of Student Evolution Acceptance—I-SEA. International Journal of Science Education, 34(11), 1637–1666. https://doi.org/10.1080/09500693.2012.702235

Nehm, R. H., & Reilly, L. (2007). Biology majors’ knowledge and misconceptions of natural selection. AIBS Bulletin, 57(3), 263–272.

Nehm, R. H., & Schonfeld, I. S. (2007). Does Increasing Biology Teacher Knowledge of Evolution and the Nature of Science Lead to Greater Preference for the Teaching of Evolution in Schools? Journal of Science Teacher Education, 18(5), 699–723. https://doi.org/10.1007/s10972-007-9062-7

Newport, F. (2007, June 11). Majority of Republicans Doubt Theory of Evolution. Retrieved January 29, 2018, from http://news.gallup.com/poll/27847/majority-republicans-doubt-theory-evolution.aspx

Owens, K., & Foos, A. (2007). A course to meet the nature of science and inquiry standards within an authentic service learning experience. Journal of Geoscience Education, 55(3), 211–217.

Ozdemir, G., & Dikici, A. (2017). Relationships between scientific process skills and scientific creativity: Mediating role of nature of science knowledge. Journal of Education in Science, Environment and Health, 3(1), 52–68.

Pew Research Center. (2015). Americans, Politics, and Science Issues (p. 175). Retrieved from http://assets.pewresearch.org/wp-content/uploads/sites/14/2015/07/2015-07-01_science-and-politics_FINAL-1.pdf

Pigliucci, M. (2008). Denying evolution: Creationism, scientism, and the nature of science. Sunderland, MA: Sinauer Associates, Inc.

Pobiner, B. (2016). Accepting, understanding, teaching, and learning (human) evolution: Obstacles and opportunities. American Journal of Physical Anthropology, 159, 232–274. https://doi.org/10.1002/ajpa.22910

R Core Team. (2017). R: A Language and Environment for Statistical Computing (Version 3.4.1). Vienna: R Foundation for Statistical Computing. Retrieved from http://www.R-project.org/

Resolution on Scientific Creationism. (1982). SBC Annual. Retrieved from http://www.sbc.net/resolutions/967

Rice, J. W., Olson, J. K., & Colbert, J. T. (2011). University Evolution Education: The Effect of Evolution Instruction on Biology Majors’ Content Knowledge, Attitude Toward Evolution, and Theistic Position. Evolution: Education and Outreach, 4(1), 137–144. https://doi.org/10.1007/s12052-010-0289-y

Richardson, J. T. E. (2011). Eta squared and partial eta squared as measures of effect size in educational research. Educational Research Review, 6(2), 135–147. https://doi.org/10.1016/j.edurev.2010.12.001

Rissler, L. J., Duncan, S. I., & Caruso, N. M. (2014). The relative importance of religion and education on university students’ views of evolution in the Deep South and state science standards across the United States. Evolution: Education and Outreach, 7(1), 24.

Romine, W. L., Walter, E. M., Bosse, E., & Todd, A. N. (2017). Understanding patterns of evolution acceptance-A new implementation of the Measure of Acceptance of the Theory of Evolution (MATE) with Midwestern university students. Journal of Research in Science Teaching, 54(5), 642–671. https://doi.org/10.1002/tea.21380

RStudio Team. (2016). RStudio: Integrated Development Environment for R (Version 1.0.153). Boston: RStudio, Inc. Retrieved from http://www.rstudio.com/

Rubba, P. A., & Andersen, H. O. (1978). Development of an instrument to assess secondary school students understanding of the nature of scientific knowledge. Science Education, 62(4), 449–458.

Rutherford, A. (2001). Introducing ANOVA and ANCOVA: A GLM approach. Thousand Oaks, CA: SAGE Publications, Inc.

Rutledge, M. L., & Mitchell, M. A. (2002). High School Biology Teachers’ Knowledge Structure, Acceptance & Teaching of Evolution. The American Biology Teacher, 64(1), 21–28. https://doi.org/10.1662/0002-7685(2002)064[0021:HSBTKS]2.0.CO;2

Rutledge, M. L., & Sadler, K. C. (2007). Reliability of the Measure of Acceptance of the Theory of Evolution (MATE) instrument with university students. The American Biology Teacher, 69(6), 332–335.

Rutledge, M. L., & Warden, M. A. (1999). The development and validation of the measure of acceptance of the theory of evolution instrument. School Science and Mathematics, 99(1), 13–18.

Shermer, M. (2006). Why Darwin Matters: The Case Against Intelligent Design. New York: Owl Books.

Short, S. D., & Hawley, P. H. (2012). Evolutionary Attitudes and Literacy Survey (EALS): Development and Validation of a Short Form. Evolution: Education and Outreach, 5(3), 419–428. https://doi.org/10.1007/s12052-012-0429-7

Sinatra, G. M., Southerland, S. A., McConaughy, F., & Demastes, J. W. (2003). Intentions and beliefs in students’ understanding and acceptance of biological evolution. Journal of Research in Science Teaching, 40(5), 510–528. https://doi.org/10.1002/tea.10087

Smith, M. U., Snyder, S. W., & Devereaux, R. S. (2016). The GAENE-Generalized Acceptance of EvolutioN Evaluation: Development of a new measure of evolution acceptance. Journal of Research in Science Teaching, 53(9), 1289–1315. https://doi.org/10.1002/tea.21328

Snyder, J. J., Sloane, J. D., Dunk, R. D. P., & Wiles, J. R. (2016). Peer-Led Team Learning Helps Minority Students Succeed. PLOS Biology, 14(3), e1002398. https://doi.org/10.1371/journal.pbio.1002398

The Clergy Letter Project. (2004). Retrieved January 24, 2018, from http://www.theclergyletterproject.org/

Trani, R. (2004). I Won’t Teach Evolution; It’s against My Religion. And Now for the Rest of the Story … The American Biology Teacher, 66(6), 419–427. https://doi.org/10.2307/4451708

Van Hiel, A., Kossowska, M., & Mervielde, I. (2000). The relationship between openness to experience and political ideology. Personality and Individual Differences, 28(4), 741–751.

Walker, J. D., Wassenberg, D., Franta, G., & Cotner, S. (2017). What Determines Student Acceptance of Politically Controversial Scientific Conclusions? Journal of College Science Teaching, 47(2), 46–56.

Walls, L. (2016). Awakening a dialogue: A critical race theory analysis of U. S. nature of science research from 1967 to 2013. Journal of Research in Science Teaching, 53(10), 1546–1570. https://doi.org/10.1002/tea.21266

Weisberg, D. S., Landrum, A. R., Metz, S. E., & Weisberg, M. (2018). No Missing Link: Knowledge Predicts Acceptance of Evolution in the United States. BioScience, 68(3), 212–222. https://doi.org/10.1093/biosci/bix161

Wiles, J. R., & Alters, B. (2011). Effects of an Educational Experience Incorporating an Inventory of Factors Potentially Influencing Student Acceptance of Biological Evolution. International Journal of Science Education, 33(18), 2559–2585. https://doi.org/10.1080/09500693.2011.565522

Woods, C. S., & Scharmann, L. C. (2001). High school students’ perceptions of evolutionary theory. Electronic Journal of Science Education, 6(2), 1–21.

